# SAGA1 and SAGA2 promote starch formation around proto-pyrenoids in Arabidopsis chloroplasts

**DOI:** 10.1101/2023.11.25.568654

**Authors:** Nicky Atkinson, Rhea Stringer, Stephen R Mitchell, David Seung, Alistair J. McCormick

## Abstract

The pyrenoid is a chloroplastic microcompartment in which most algae and some terrestrial plants condense the primary carboxylase, Rubisco (ribulose-1,5-bisphosphate carboxylase/oxygenase) as part of a CO_2_-concentrating mechanism that improves the efficiency of CO_2_ capture. Engineering a pyrenoid-based CO_2_-concentrating mechanism (pCCM) into C3 crop plants is a promising strategy to enhance yield capacities and resilience to the changing climate. Many pyrenoids are characterized by a sheath of starch plates that is proposed to act as a barrier to limit CO_2_ diffusion. Recently, we have reconstituted a phase-separated ‘proto-pyrenoid’ Rubisco matrix in the model C3 plant *Arabidopsis thaliana* using proteins from the alga with the most well studied pyrenoid, *Chlamydomonas reinhardtii* (1). Here we describe the impact of introducing the Chlamydomonas proteins StArch Granules Abnormal 1 (SAGA1) and SAGA2, which are associated with the regulation of pyrenoid starch biogenesis and morphology. We show that SAGA1 localizes to the proto-pyrenoid in engineered Arabidopsis plants, which results in the formation of atypical spherical starch granules enclosed within the proto-pyrenoid condensate and adjacent plate-like granules that partially cover the condensate, but without modifying the total amount of chloroplastic starch accrued. Additional expression of SAGA2 further increases the proportion of starch synthesised as adjacent plate-like granules that fully encircle the proto-pyrenoid. Our findings pave the way to assembling a diffusion barrier as part of a functional pCCM in vascular plants, whilst also advancing our understanding of the roles of SAGA1 and SAGA2 in starch sheath formation and opening novel avenues for engineering starch morphology.

## Introduction

Almost all life on Earth is fuelled by the carboxylase activity of ribulose-1,5-biphosphate carboxylase oxygenase (Rubisco), the primary enzyme used to capture CO_2_ for growth and metabolism. Nevertheless, Rubisco has a relatively slow turnover rate and a competitive oxygenase reaction, which can lead to energy and CO_2_ losses (2). Most plants (i.e. C3 plants) rely on passive diffusion of ambient CO_2_ into the chloroplast where Rubisco catalyses CO_2_ uptake. However, Rubisco is typically only half-saturated at CO_2_ concentrations inside chloroplasts of C3 plants, which leads to a significant protein investment in the Rubisco pool but still allows for counter-productive oxygenation reactions that can occupy up to a third of the active sites of Rubisco (3). In C3 staple crops, such as wheat and rice, photorespiration causes significant yield losses (4). Several recent lab- and field-based studies have shown that improving the efficiency of CO_2_ capture by Rubisco is a promising strategy to increase crop yields (5–7). Such research is important to ameliorate global food insecurity concerns (8), particularly as rising temperatures due to climate change will worsen the impacts of photorespiration despite increasing CO_2_ levels (9, 10).

To overcome the limitations of Rubisco, many important lineages of algae and some non-vascular terrestrial plants (hornworts) have evolved a pyrenoid-based CO_2_-concentrating mechanism (pCCM) to elevate the level of CO_2_ around Rubisco (11). The pyrenoid is a membrane-less condensate formed by the liquid-liquid phase separation of Rubisco into a matrix that is fed with above ambient concentrations of CO_2_, which preferentially drives the Rubisco carboxylation reaction and increases its turnover rate (12). Pyrenoids are typically characterised by single or multiple traversing thylakoid membranes that likely supply CO_2_ to the Rubisco matrix and a sheath of curved starch plates that surround the matrix (11). The starch sheath is predicted to act as a diffusion barrier to limit CO_2_ diffusion, and could also restrict inward diffusion of O_2_ to prevent photorespiration (13, 14). Modelling predicts that reconstituting a functional pCCM into C3 crop plants has the potential to significantly enhance crop yield potentials (14, 15).

The green alga *Chlamydomonas reinhardtii* (hereafter Chlamydomonas) has the most well characterised pCCM (for recent reviews see 15–17). In Chlamydomonas, Rubisco condensation in the pyrenoid is facilitated by weak multivalent interactions between the small subunit of Rubisco (RbcS) and the disordered linker protein EPYC1 (Cre10.g436550) via five Rubisco binding motifs (RBMs) on EPYC1 (18–20). RBMs are thought to play a key role in pyrenoid assembly as they are found in many pyrenoid-localized proteins, including the starch sheath-associated proteins SAGA1 (StArch Granules Abnormal 1, Cre11.g467712) and SAGA2 (Cre09.g394621) (21, 22). SAGA1 is a ∼180-kDa protein with two RBMs and a predicted starch-binding domain (Carbohydrate Binding Module 20 - CBM20) located near the C-terminus. SAGA1 localizes to puncta at the periphery of the Rubisco matrix at the interface between the matrix and the starch sheath and is required for normal starch sheath formation and pyrenoid morphology, although the underlying molecular mechanism is unknown (21). Cells lacking SAGA1 have thin and elongated pyrenoid starch granules and are characterised by multiple pyrenoids that lack traversing thylakoid membranes. SAGA2 is a ∼190-kDa protein that is 30% identical to SAGA1, and has four RBMs and a CBM20 domain (22, 23). Like SAGA1, SAGA2 has a predicted C-terminal starch-binding domain and localizes to the matrix-starch sheath interface, although SAGA2 appears to cover the surface of the matrix more homogeneously than SAGA1. These observations together with previous work (24–26) support a model where SAGA1 and SAGA2 mediate adherence of the starch sheath to the matrix.

The location and shape of the starch sheath that surrounds the pyrenoid in Chlamydomonas differs from that of the transitory starch granules used for carbohydrate storage (27). In both green algae and plant leaf mesophyll cells, transitory starch granules are discrete and occupy ‘stromal pockets’ between thylakoid membranes (28). Recent work in Arabidopsis has shown that newly initiated starch granules can initially coalesce and then undergo anisotropic growth to form characteristic lenticular granules (29). Several proteins implicated in granule initiation are thylakoid-associated (30–32), indicating an important role for the thylakoid membranes in transitory starch biosynthesis (33). Given their distinct shape and placement, the components and mechanisms of starch sheath formation around pyrenoids likely differ from those for starch formation in the stroma. For example, SAGA1 and SAGA2 could regulate the sites of starch sheath formation around the pyrenoid via recruitment of starch-forming enzymes from the stroma and/or those associated with the thylakoid membrane (34).

Significant progress has been made towards building a functional pCCM in a C3 land plant (1, 26, 35, 36), including the reconstitution of a phase-separated Rubisco matrix in the model C3 plant *Arabidopsis thaliana* (hereafter Arabidopsis). Atkinson et al. (1) recently demonstrated in Arabidopsis with reduced native Rubisco small subunit (AtRbcS) levels that expression of EPYC1 and a Chlamydomonas RbcS (CrRbcS) was sufficient to condense Rubisco into a single ‘proto-pyrenoid’ in each chloroplast. Modelling of the Chlamydomonas pCCM predicts that the generation of a starch sheath around the condensate will be important for improving the energetic efficiency of the pyrenoid by i) decreasing wasteful CO_2_ leakage, and ii) improving Rubisco activity by increasing the concentration of CO_2_ in the Rubisco matrix up to 10-fold (14). Here, we examined whether SAGA1 and SAGA2 could recruit starch to the proto-pyrenoid. Remarkably, we observed the formation of atypical spherical starch granules enclosed within the condensate, and the recruitment of up to 71% of total chloroplastic starch within and around the condensate. Subsequent serial block-face scanning electron microscopy imaging revealed the formation of plate-like starch granules that encircled the condensate in plants expressing SAGA1 and SAGA2, indicating that SAGA1 and SAGA2 can regulate the recruitment and shaping of starch around the proto-pyrenoid *in planta*.

## Results

### SAGA1 localizes around the proto-pyrenoid in Arabidopsis

To determine the expression pattern of SAGA1 when introduced into Arabidopsis, we stably expressed SAGA1 fused to the mCherry fluorescent protein (SAGA1::mCherry) in the S2_Cr_ background. S2_Cr_ is a *1a3b* RbcS mutant complemented with CrRbcS2 (Cre02.g120150) that produces a functional hybrid Rubisco (AtRBcL/CrRbcS2) with catalytic properties comparable to WT Rubisco (36). S2_Cr_ lacks the EPYC1 linker protein required for Rubisco condensation (**Fig. 1*A***). Confocal microscopy imaging of leaves from the resulting transgenic plants (S1) showed that SAGA1::mCherry was located in the chloroplast, demonstrating that the native SAGA1 transit peptide was sufficient to target SAGA1 to the chloroplast in plants despite the SAGA proteins having no homology with any known land plant proteins, according to BLAST analyses. SAGA1::mCherry appeared to form fluorescent puncta within each chloroplast (**Fig. 1*B***). However, transmission electron microscopy (TEM) images of S1 chloroplasts showed no apparent structural differences to those in the S2_Cr_ background (**Fig. 1*C***), indicating that the SAGA1::mCherry puncta were likely soluble.

**Figure 1.**
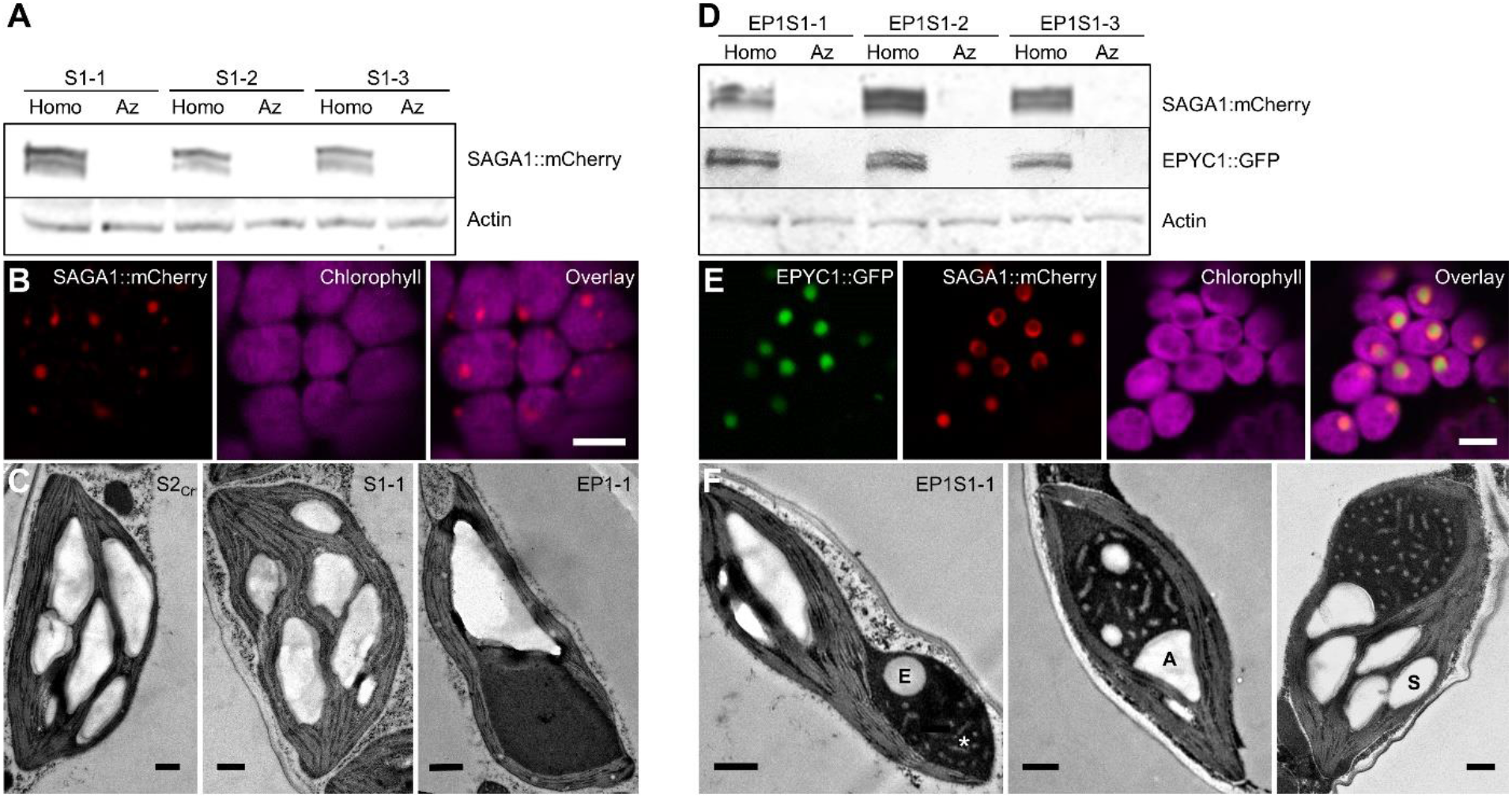
Expression of SAGA1 leads to the formation of atypical starch granules associated with the proto-pyrenoid and a novel condensate phenotype. **A.** Immunoblots showing expression of SAGA1::mCherry (S1) in three independent T3 homozygous (homo) lines and respective azygous (Az) segregants in the S2_Cr_ background (see **Fig. S13** for uncropped immunoblots). **B.** SAGA1::mCherry localises to the chloroplast in S2_Cr_. **C.** TEM of chloroplasts from S2_Cr_, S1-1, and EPYC1::GFP in S2_Cr_ (EP1-1) (Atkinson et al., 2020) **D.** Immunoblots showing expression of SAGA1::mCherry and EPYC1::GFP (EP1S1) in three independent T3 homozygous lines and respective azygous segregants in the S2_Cr_ background. **E.** SAGA1::mCherry localises to the proto-pyrenoid and forms a ring around EPYC1::GFP. **F.** Representative TEM images of EP1S1-1 chloroplasts with atypical spherical enclosed (E), adjacent (A) and typical stromal (S) starch granule labelled. The lighter-staining pattern is highlighted with an asterisk. Scale bar = 0.5 µm.

Next, we stably co-expressed SAGA1::mCherry with EPYC1::GFP in the S2_Cr_ background and generated three independent EP1S1 transgenic plant lines (**Fig. 1*D***). As observed previously (1), a single proto-pyrenoid condensate formed within each chloroplast that displaced the thylakoid membranes (**Fig. 1*E***). The fluorescence signal for SAGA1::mCherry was clearly visible in the vicinity of the condensate, but rather than co-localizing, SAGA1::mCherry preferentially formed a ring around the condensate periphery. This was consistent with the localization pattern observed in Chlamydomonas, where SAGA1 surrounds the pyrenoid presumably forming an interface between the Rubisco matrix and the starch sheath (21, 22). Although the ring localization pattern was frequently observed in all three EP1S1 lines, in some chloroplasts SAGA1::mCherry appeared more dispersed throughout the condensate (**Fig. S1*A***). TEM imaging of EP1S1 condensates revealed two notable differences compared to those described previously with only EPYC1 (EP1-1) (**Fig. 1*C***) (1). First, EP1S1 condensates contained a lighter-staining pattern spread throughout the matrix (**Fig. 1*F***). Depending on the cross-section, these patterns were observed as puncta or tubule-like structures within or around the edge of the matrix, and sometimes as concentric rings (**Fig. S1*B***). Second, we observed round starch granules that appeared to be enclosed within the EP1S1 condensate. In addition, starch granules with an atypical angular shape were often observed adjacent to the condensate, in which the sides of granules in contact with the condensate were characterised by a rounded surface.

### EP1S1 condensates retain liquid-like properties and recruit transient starch

Chlamydomonas pyrenoids behave as a liquid-like phase-separated compartment (37–39), a phenomenon that was observed previously for proto-pyrenoid condensates in Arabidopsis (1). To investigate the impact of the addition of SAGA1, we first isolated condensates from EP1S1 by gentle centrifugation (16), and confirmed that they were enriched in SAGA1, EPYC1, the native large subunit of Rubisco (AtRbcL; AtCg00490) and CrRbcS2 (**Fig. 2*A*** **and** ***B***). AtRbcS (the lower band representing AtRbcS1B and AtRbcS2B (1)) appeared absent, indicating that the Rubisco matrix in the EP1S1 condensate consisted of hybrid Rubisco. In line with previous work (1), isolated condensates from S2_Cr_ plants expressing only EPYC1 (EP1) showed enrichment of EPYC1 and hybrid Rubisco in the condensate. In contrast, S2_Cr_ plants expressing only SAGA1 (S1) showed no enrichment of Rubisco or SAGA1 in the pellet fraction, which was consistent with the observed absence of a condensate in these lines (**Fig. 1*C***). To investigate the liquid-like behaviour of EP1S1 condensates, we then performed fluorescence recovery after photobleaching (FRAP) assays. The rate of full recovery from photobleaching for EP1S1 condensates was similar to that for EP1 condensates (i.e. ∼20 s) and the half FRAP (*T*_0.5_) values were comparable between datasets, indicating that SAGA1 does not affect liquid-like mixing (**Fig. 2*C***).

**Figure 2.**
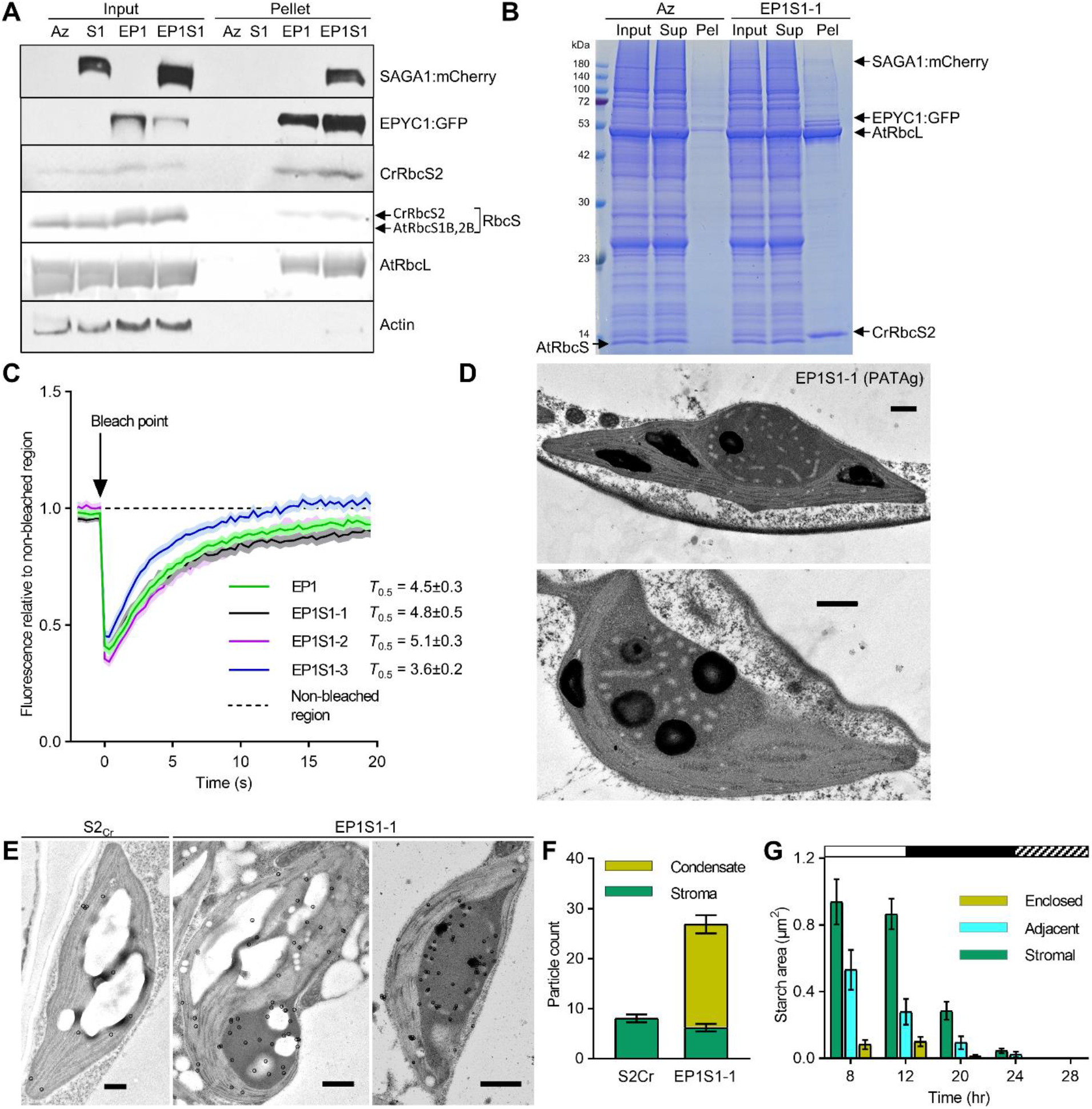
Proto-pyrenoids in EP1S1 plants contain SAGA1 and EPYC1, are liquid-like and contribute to the formation of atypical starch granules. **A.** SAGA1, EPYC1 and Rubisco protein levels in S2_Cr_ T3 homozygous lines expressing SAGA1 (S1; S1-1), EPYC1 (EP1), SAGA1 and EPYC1 (EP1S1; EP1S1-1) and an azygous EP1S1-1 segregant (Az) are shown. Whole leaf tissue samples (input) and pelleted condensate extracts (pellet) were assessed by immunoblot analyses with anti-SAGA1, anti-EPYC1, anti-CrRbcS2 or polyclonal anti-Rubisco (AtRbcL, AtRbcS1B (At5g38430), AtRbcS2B (At5g38420) and CrRbcS2 are shown) antibodies. Anti-actin is shown as an input loading control. Molecular weights: AtRbcL, 55 kDa; AtRbcS1B/2B, 14.8 kDa; CrRbcS2, 15.5 kDa; EPYC1, 34 kDa; SAGA1, 180 kDa. The immunoblots shown were derived from the same experiment and gels/blots were processed in parallel. **B.** Coomassie-stained SDS-PAGE gel showing the composition of the input, the supernatant following condensate extraction and centrifugation (Sup) and the pelleted condensate (Pel). **C.** Fluorescence recovery assays of bleached areas for condensates in EP1 and EP1S1-1, -2 and -3 lines compared to non-bleached regions. Each dataset is the mean ± SEM of 17-23 individual condensates from different chloroplasts. Half FRAP (*T*_0.5_) values (the timepoint when condensates achieve 50% bleaching recovery) are shown for each dataset. **D.** Example TEM images of periodic acid-thiocarbohydrazide-silver proteinate (PATAg) stained sections from EP1S1-1. **E.** Representative immunogold labelling of SAGA1 in EP1S1-1. Gold nano-particles are highlighted with circles. Scale bars for all TEM images = 0.5 µm. **F.** Average number of gold particles observed in the condensate versus the stroma. Mean ± SEM, n = 23 (S2_Cr_) and 48 (EP1S1-1). **G.** Starch area in EP1S1-1 plants based on TEM images taken over a 12:12 hr light: dark cycle followed by a 4 hr in the dark (see **Fig. S3** for more details). White bar = light, black bar = dark, shaded bar = starvation in dark. Starch granules were scored as stromal, adjacent to the condensate (i.e. between 10 and 90 % of the starch granule periphery was in contact with the condensate) or enclosed by the condensate (i.e. greater than 90 % of the granule periphery was in contact with the condensate). Each timepoint represents the mean ± SEM of 22-39 chloroplasts.

We next investigated the novel patterning and starch accumulation observed within the EP1S1 condensates by TEM. Periodic acid-thiocarbohydrazide-silver proteinate (PATAg) staining confirmed that the round structures within the condensates were starch granules (**Fig. 2*D***). In contrast, the lighter-staining patterns were not stained by PATAg, and thus were unlikely to be glucan-based. The patterned structures were also not stained during osmication, indicating that they are likely not comprised of lipid, and therefore not membrane-based **(Fig. S2).** Immunogold analysis demonstrated an enrichment of SAGA1 around the round starch granules and in the vicinity of the patterned structures (**Fig. 2*E* and 2*F***), indicating that SAGA1 was directly associated with both features. To compare the turnover of stromal and condensate-associated starch, we then visually tracked the area of starch granules in EP1S1 chloroplasts by TEM over a diel light-dark cycle and a subsequent 4 hr starvation period in the dark (**Fig. 2*G*** and **Fig. S3*A***). As expected, starch levels were highest at 8 and 12 hr during the light period, with 73% of starch located in the stroma, 18% adjacent to the condensate and 10% enclosed by the condensate at the end of a 12 hr day. During the dark period the turnover rates for adjacent and enclosed granules were 3- and 8-fold slower compared to stromal starch (**Fig. S3*B***). But interestingly, we observed that the proportional turnover rates for all starch granules were similar, such that both stromal and condensate-associated starch were degraded by the perceived end of the night (24 hr) (**Fig. S3*C***) (40). Overall, in EP1S1, the adjacent and enclosed starch granules could be turned over in the night, despite their distinctive shape and localisation away from the stromal pockets (29).

### SAGA2 localises to the condensate and around starch granules

SAGA2 is a strong candidate for regulating the formation of the starch sheath in Chlamydomonas based on similarities in protein sequence to SAGA1 and comparable localisation to the pyrenoid starch-matrix interface (22). We first stably expressed SAGA2 fused to mNeon in the Arabidopsis background S2_Cr_. Similar to SAGA1, the native SAGA2 transit peptide could target SAGA2::mNeon to the chloroplast stroma, where it appeared to form rings or curved patterns (**Fig. 3*A***). We hypothesised that the predicted starch-binding domain of SAGA2 might be causing localisation to the edges of starch granules (41). We then expressed EPYC1::GFP in the S2_Cr_ background with SAGA2 fused to an mCherry tag to generate the transgenic plant line EP1S2. Unfortunately, we were unable to examine EP1S2 plants past the T1 generation due to apparent silencing of SAGA2 in the T2 generation. Proto-pyrenoid condensates formed as expected in T1 lines, and SAGA2::mCherry co-localised with EPYC1::GFP throughout the condensates, which were characterised by a crescent or concave shape (**Fig. 3*B***). SAGA2 was also observed to localise in circles within and away from the condensate (**Fig. 3*B***, white and red arrows), presumably at the edges of starch granules enclosed within the condensate and in the stroma, respectively. When viewed by TEM, starch granules could be observed within the EP1S2 condensates (**Fig. 3*C***). In contrast to SAGA1, no lighter-staining patterns were found in the condensate when expressing SAGA2. Together, our imaging data suggested that SAGA2 interacts differently with the condensate compared to SAGA1 but is also able to recruit starch to the matrix.

**Figure 3.**
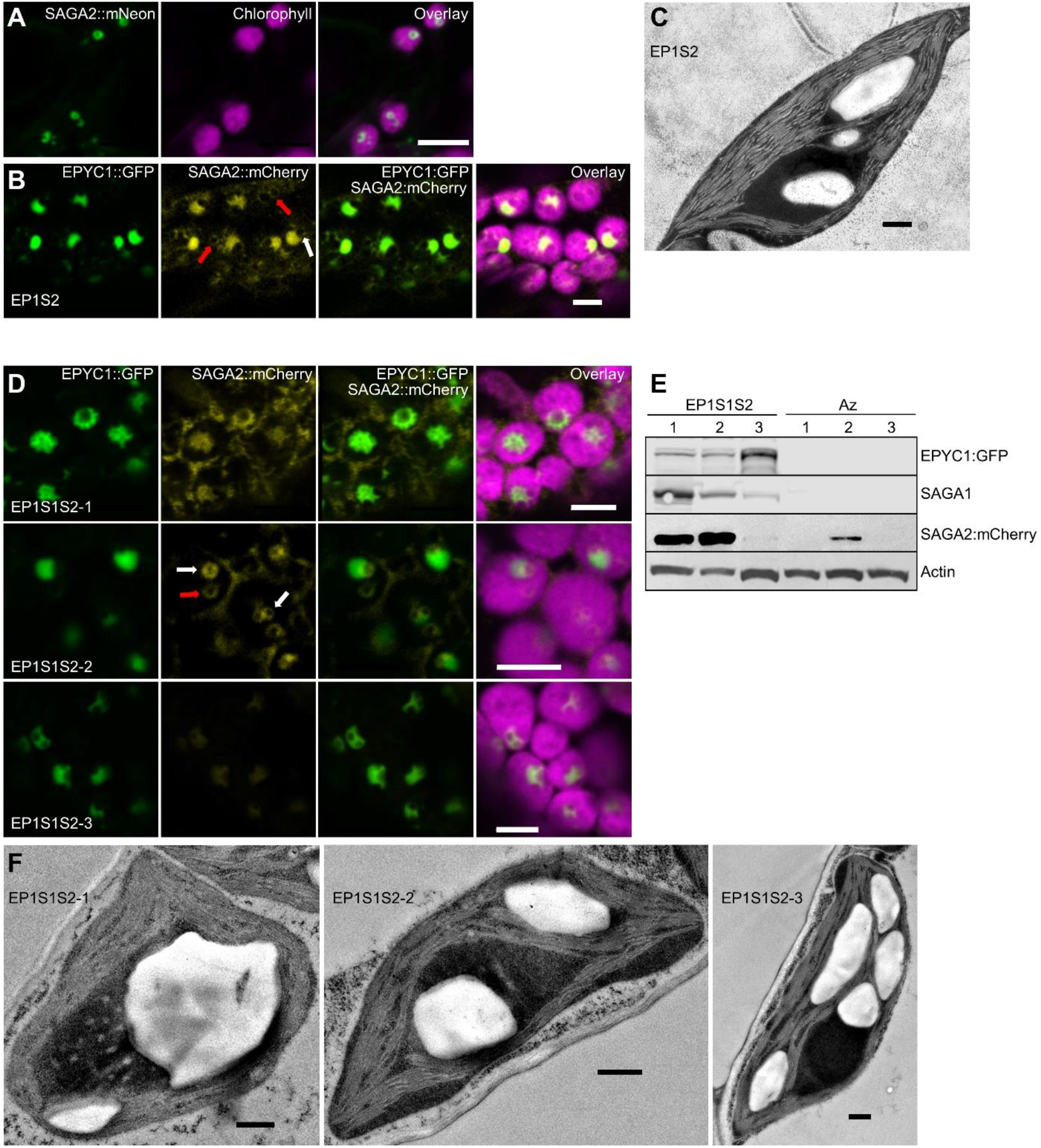
Proto-pyrenoids in plants expressing SAGA1, SAGA2 and EPYC1 are associated with the formation of large adjacent starch granules. **A.** SAGA2::mNeon localises to the chloroplast in S2_Cr_. **B.** SAGA2::mCherry mainly co-localises with EPYC1::GFP in condensates in S2_Cr_ (EP1S2). White and red arrows illustrate examples of SAGA2 encircling a cavity (likely a starch granule) within and away from a condensate, respectively. **C.** Representative TEM image of EP1S2. **D.** SAGA2::mCherry mainly co-localises to the distorted condensates when expressed with SAGA1 and EPYC1::GFP in S2_Cr_ (EP1S1S2-1, -2 and -3). **E.** Immunoblots showing differential expression of SAGA1, SAGA2 and EPYC1 in three independent T3 EP1S1S2 lines and their azygous segregants (note: Az2 still showed SAGA2 expression and was excluded from further analyses). **F.** Representative TEM images of each EP1S1S2 lines. Scale bar on all confocal and TEM images = 5 µm.

We next stably expressed both untagged SAGA1 and SAGA2::mCherry with EPYC1::GFP in the S2_Cr_ background to investigate if co-expression might further increase starch granule association with the proto-pyrenoid condensate. Three independent EP1S1S2 lines were generated that expressed all three transgenes (**Fig. 3*D* and 3*E***). Confocal microscopy imaging showed that the condensates in EP1S1S2 lines were distorted, with one or several cavities visible within or at the edges of the condensate, consistent with the presence of starch granules. Notably, in the line with highest SAGA2 expression (EP1S1S2-2), SAGA2::mCherry appeared to form circles around the cavities within the condensates (**Fig. 3*D***, white arrows), and circles not associated with the condensate (red arrow), supporting our earlier observation that SAGA2 can also bind starch granules outside the condensate (**Fig. 3*B***). To better visualise this phenomenon, we used three unique fluorescent tags to generate an additional S2_Cr_ line expressing SAGA1::mCherry, SAGA2::BFP and EPYC1:GFP. Consistent with other EP1S1S2 lines, the condensates were distorted, SAGA1::mCherry localised primarily with the condensate, whereas SAGA2::BFP localised to a circle at the edge of the condensate (**Fig. S4**). Subsequent TEM images confirmed that the condensate distortions in the EP1S1S2 lines were due to the presence of one or several large starch granules within or associated with the matrix (**Fig. 3*F*** **and S5**). The highest expressing SAGA1 line (EP1S1S2-1) was often characterised by a large circular starch granule within the condensate and retained the lighter-staining patterns in the matrix observed in EP1S1 chloroplasts (**Fig. 1 and 2**). In EP1S1S2-2 the ratio of SAGA1:SAGA2 expression was decreased compared to EP1S1S2-1, the starch granules associated with the condensate were relatively smaller, and the patterned structures in the matrix were absent. Some chloroplasts in EP1S1S2-2 appeared to contain two condensates, each with enclosed starch granules (**Fig. S5** and **S*6A***). EP1S1S2-3 was characterised by high expression levels for EPYC1 but relatively little SAGA1 and SAGA2, and much less starch associated with the condensate (**Fig. S6*B***).

### SAGA2 interacts with the α-helical repeat sequences of EPYC1

A previous protein interactome study has suggested that SAGA2 can interact with both EPYC1 and Rubisco (23). Here, we used yeast two-hybrid assays to further understand these putative interactions (**Fig. S*7***). Similar to SAGA1, SAGA2 interacted with both CrRbcS and the Chlamydomonas large subunit of Rubisco (CrRbcL), albeit in the absence of chaperones (**Fig. S7*B***) (21). It remains unclear how SAGA1 and SAGA2 might interact with CrRbcL. SAGA2 showed a relatively strong interaction with EPYC1, similar in strength to that observed between EPYC1 and CrRbcS (26). To investigate how EPYC1 interacts with SAGA2, we then assayed SAGA2 with an EPYC1 helix KO mutant, wherein the repeated α-helix sequences ‘K/WQELESL’ were each replaced by a string of seven alanine residues (**Fig. S7*C***). SAGA2 could not interact with the EPYC1 helix KO, indicating that EPYC1 interacts with SAGA2 through the same α-helices sequences required for interaction with CrRbcS (19). We subsequently screened SAGA2 with a single EPYC1 α-helix repeat sequence with substitutions in residues known to interact with CrRbcS. Replacement of the negatively charged glutamic acid residues E66 (critical for interaction with CrRbcS) and E68 with valine (V) did not reduce the interaction between EPYC1 and SAGA2. However, replacement of the hydrophobic leucine residues L67 and L70 with positively charged arginine (R) abolished the interaction. The impact of substituting either L67 or L70 with R was minor, but replacement of L70 with negatively charged E or histidine (H) abolished or strongly reduced the interaction, respectively. Thus, it appears that the hydrophobic residues in the EPYC1 α-helix are potentially more important for SAGA2 interaction than those that mediate salt-bridge interactions with CrRbcS (**Fig. S7E**) (19). To investigate where SAGA2 interacts with EPYC1, the SAGA2 sequence was divided into five regions based on predicted areas of coiled coil domains and disorder (42) (**Fig S7*D*** and **S7*F***). Only one region containing several coiled coil domains (region 3) showed an interaction with EPYC1, which was relatively weak compared to the full length SAGA2. When region 3 was subdivided into three smaller sequences (3A-C), 3B showed only a minimal interaction, suggesting that the folded structure of SAGA2, rather than one particular domain, is important for interaction with EPYC1.

### SAGA1 and SAGA2 recruit starch with distinct morphologies to the condensate

The enlarged starch granules observed in the EP1S1S2 lines prompted the question of whether SAGA1 and/or SAGA2 might increase the abundance of total starch per chloroplast. Thus, we measured the starch content in our transgenic lines grown in long days (16 hr light: 8 hr dark) at 1 hr (dawn), 8 hr (noon) and 16 hr (dusk) during the light period (**Fig. 4*A***). Despite the starch granules varying greatly in distribution and morphology between the lines (**Fig. 1-3**), we observed no consistent differences in overall starch content at any time point between lines, although EP1S1-1 samples showed a wide variation in starch content at 8 hr.

**Figure 4.**
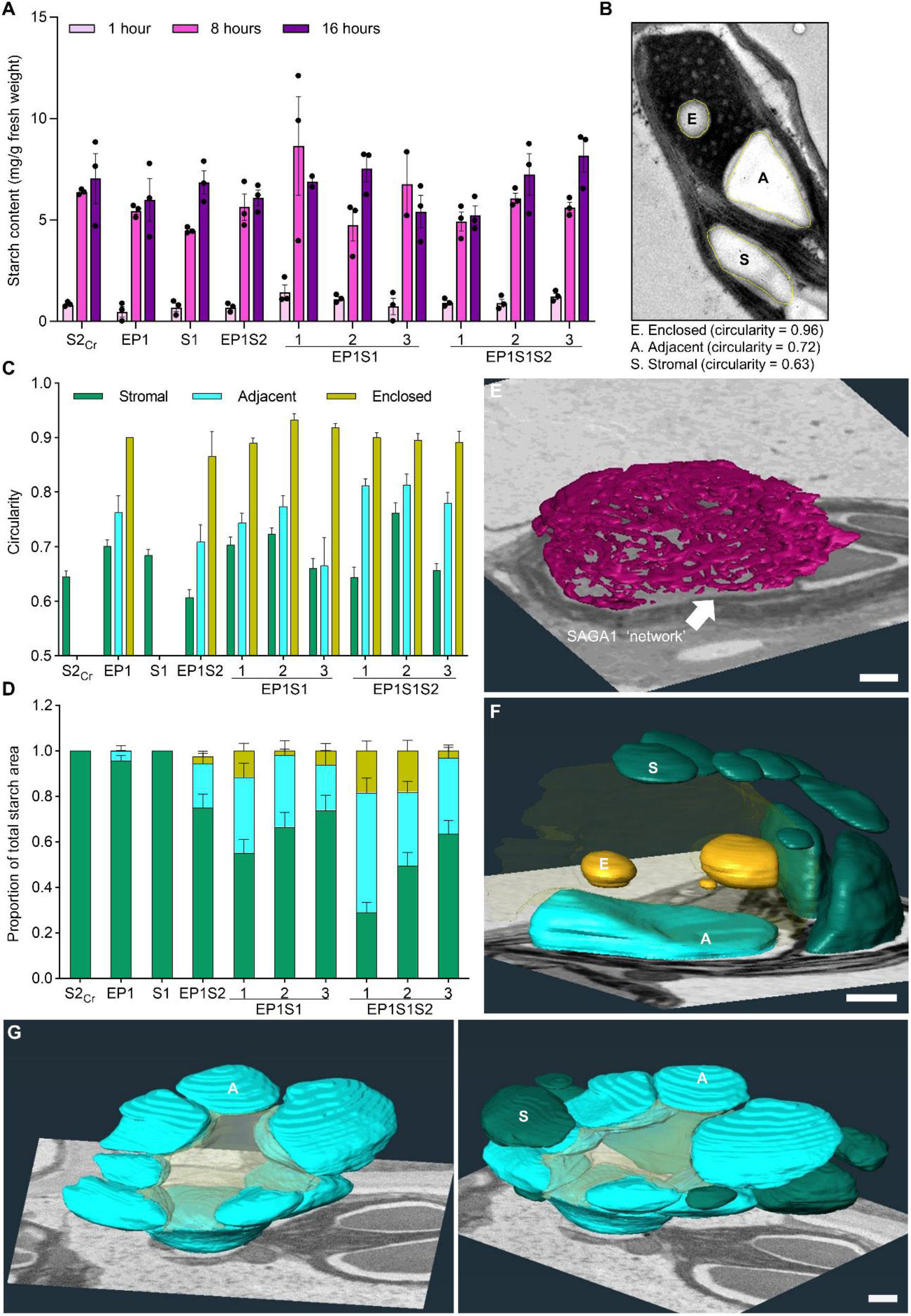
Plants expressing SAGA1, SAGA2 and EPYC1 produce plate-like starch granules that encircle the proto-pyrenoid. **A.** Leaf starch content at 1 hr, 8 hr and 16 hr into the photoperiod. Bars represent the mean ± SEM from three individual rosettes. **B.** Example of enclosed, adjacent and stromal starch granules (in EP1S1). The circularity values for each granule were calculated using the freehand selection tool in Fiji (1 = perfect circle and 0 = straight line). **C.** Circularity of starch granules for each plant line measured from TEM images. **D.** Proportion of starch area based on granules categorised as enclosed, adjacent or stromal. Starch granules were designated ‘adjacent’ if between 0-90% of perimeter bordered the condensate, and ‘enclosed’ if >90% of perimeter bordered the condensate. The bars represent the mean ± SEM of starch granules from 28-48 chloroplasts in plants imaged at the end of the photoperiod. **E.** 3-D reconstructions of the SAGA1 network in EP1S1 obtained using SBF-SEM. **F.** 3-D reconstructions of stromal (S), enclosed (E) and adjacent (A) starch granules in EP1S1. The condensate is shown in transparent yellow. **G.** 3-D reconstructions of adjacent starch granules in EP1S1S2 with (right) and without (left) stromal starch granules shown. In **F** and **G** the condensate is shown in transparent yellow. Scale bar on all SBF-SEM images = 1 µm. See **Fig. S8** for raw video data and 3-D reconstruction videos.

To further characterise the starch granules recruited to the condensates in different lines, we used TEM to quantify the shape (circularity) and area of visible starch per chloroplast. Circularity was assessed using Fiji (ImageJ), where a value of 1 represents a perfect circle (**Fig. 4*B***). The spherical starch granules enclosed by the condensate (defined by having >90% of their perimeter in contact with the condensate) achieved high circularity values of ∼0.9. In contrast, the adjacent granules (defined by having 10-90% of their perimeter in contact with the condensate) had an average circularity value of 0.72, while the typical lenticular shape of stromal starch granules gave an average value of 0.63.

Guided by these thresholds, we quantified the area of starch for enclosed, adjacent and stromal granules in different plant lines, as well as the number of starch granules in each category (**Fig. 4*C*** and **S6*C***). In EP1 plants the majority of starch granules were located in the stroma, with negligible starch adjacent to the condensate. However, expression of SAGA1 or SAGA2 led to a distinct increase in the proportion of condensate-associated starch (either enclosed or adjacent), ranging from 22.4% in our single EP1S2 line to 45% in EP1S1-1. Co-expression of SAGA1 and SAGA2 resulted in up to 71% of total starch associated with the condensate in EP1S1S2-1 (**Fig. 4D**). For the latter line, categorisation of ‘adjacent’ or ‘enclosed’ granules was challenging as the granules were often large, obscured the condensate, and made contact with the surrounding thylakoids (**Fig S*5***). Notably, the stromal starch granules in the EP1S1S2 lines were both smaller and less numerous than in other lines. Consistent with starch content measurements (**Fig. 4A**), the total starch granule number per chloroplast was generally consistent between lines with the exception of EP1S1-1, which had a slightly higher average (**Fig. S6*D***). The expression of SAGA1 correlated with the levels of condensate-associated starch in the EP1S1S2 lines despite EP1S1S2-1 and EP1S1S2-2 having similar levels of SAGA2 (**Fig. 3*E***), suggesting that SAGA1 recruits more starch to the condensate than SAGA2. Both SAGA1 and SAGA2 expression levels were much reduced in EP1S1S2-3, which had similar levels of starch association to the EP1S1 lines.

We next used serial block-face scanning electron microscopy to image and generate 3D reconstructions of the morphological features of EP1S1-1 and EP1S1S2-1. We first reconstructed the lighter-staining regions observed in condensates in EP1S1 lines (**Fig. 1F**). Based on our immunogold analysis (**Fig. 2*E***), these regions appear to contain SAGA1 protein, which could be connected via interactions with the Rubisco matrix. Remarkably, we observed that the lighter-staining regions formed an interconnected network within the condensate (**Fig. 4*E***, **Fig S8*A*** and **S8*B***). The network bears some resemblance to the system of thylakoid membranes that traverse the Chlamydomonas pyrenoid (24), although the network did not make contact with the surrounding thylakoids and did not appear to be lipid-based (**Fig. S2**). This observation raises the possibility that SAGA1 could play an additional role apart from that in starch sheath formation, perhaps as a protein scaffold to guide the traversing thylakoids in Chlamydomonas. Alternatively, it could be a SAGA1 overexpression artifact, as SAGA1 expression was driven by a strong promoter in Arabidopsis and is thus likely at higher levels than that in Chlamydomonas (15, 43), and SAGA1 was not observed to localize inside pyrenoids (21). We also observed enclosed, adjacent and stromal starch granules in EP1S1-1 (**Fig. 4*F***, **Fig S8*C*** and **S8*D***). As expected, the enclosed round starch granules were spherical and contained within the condensate. In contrast, the adjacent granules proved larger than expected and typically extended around the side of the condensate, forming a curved shape similar to the plate-like starch that constitute the pyrenoid starch sheath in Chlamydomonas. In EP1S1S2-1 several adjacent starch granules formed a ring of starch granules that encircled the condensate (**Fig. 4*G***, **Fig. S8*E*** and **S8*F***). In both EP1S1-1 and EP1S1S2-1, the stromal starch granules away from the condensate retained their characteristic lenticular shape. Consistent with our TEM data (**Fig. 1C**), condensates in EP1 plants did not associate with starch (**Fig. S8*G*** and **S8*H***), and like S1-1 and S2_Cr_ plants, produced lenticular starch granules in the stroma (**Fig. S8*I-L***).

### Expression of SAGA1 and SAGA2 does not impact photosynthesis

We analysed the growth and photosynthetic performance of three independent T3 lines for S1, EP1S1 or EP1S1S2. No consistent differences in growth were observed between S1 and EP1S1 plants and their segregant controls (**Fig. S9**). This is consistent with previous work where no differences in growth were observed between EP1 plants and segregant lines (1). However, a reduction in growth was observed for the three EP1S1S2 lines (**Fig. 5*A*** and **5*B***). To determine if the EP1S1S2 growth phenotype was due to differences in photosynthetic performance, we examined the response of CO_2_ assimilation to intercellular CO_2_ concentration under saturating light (*A*:*C*i curves) (**Fig 5*C***). The photosynthetic parameters derived from the *A*:*C*i curves were comparable between plants measured under photorespiratory (ambient O_2_) or non-photorespiratory (2% O_2_) conditions (**Fig. S10** and **S11**), which indicated that the observed growth phenotype was not due to reduced Rubisco activity. This is consistent with previous work showing that Rubisco condensation *in planta* does not appear to impact Rubisco activity (1), and suggests that the recruitment of starch plate-like granules around the condensate is not impeding CO_2_ diffusion in EP1S1S2 lines.

**Figure 5.**
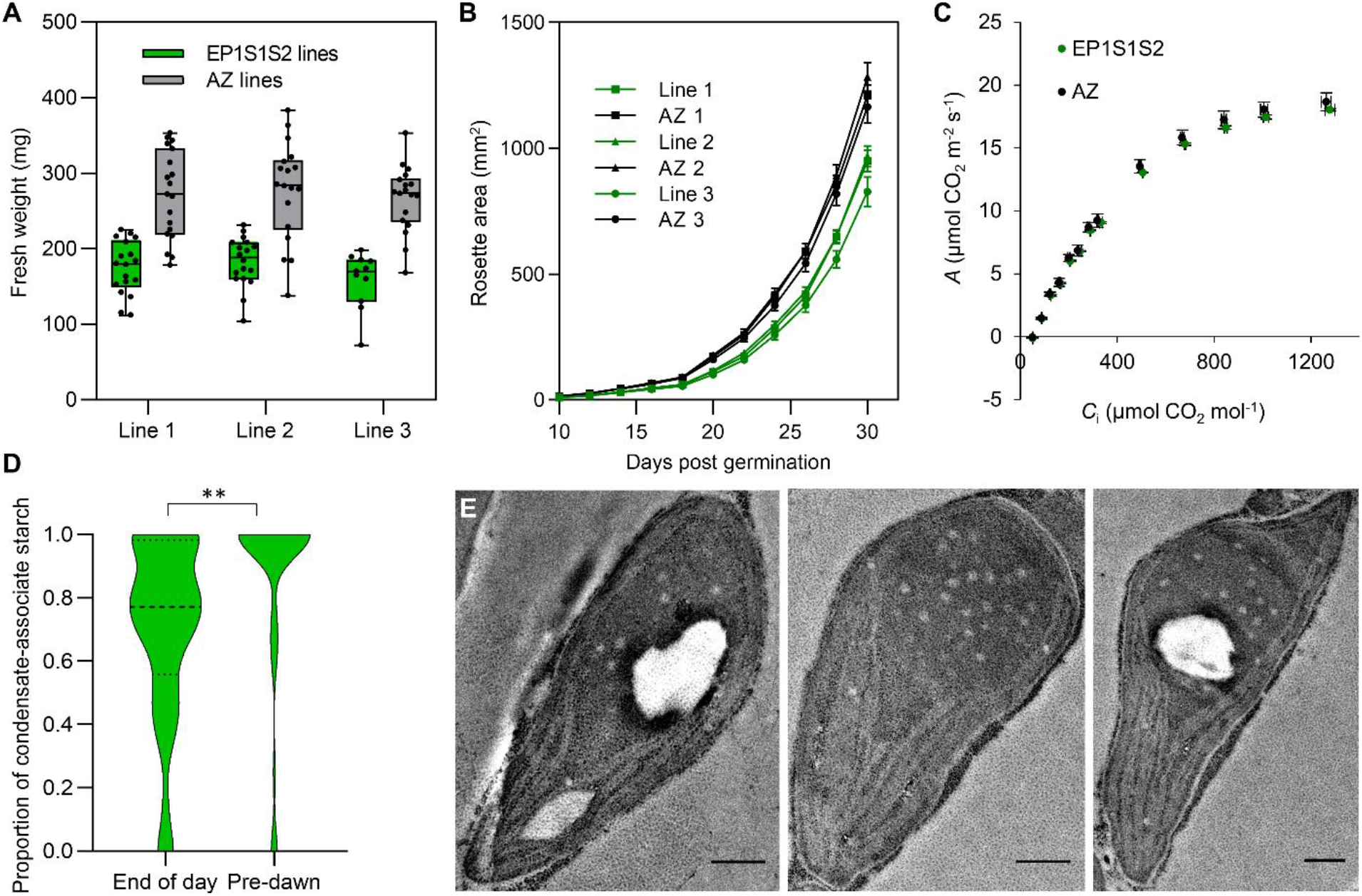
Growth is reduced but not photosynthetic performance in EP1S1S2 lines. **A.** Fresh weight of EP1S1S2 lines and their azygous (Az) segregants 32 days after germination. The S2_Cr_ background line was used as a control for line 1 as the Az segregant line had consistent germination and growth problems. **B.** Rosette expansion for S1 and EP1S1 lines measured over 30 days post germination. Error bars show the mean ± SEM of 12-21 individual rosettes. **C.** Net CO_2_ assimilation (*A*) based on sub-stomatal [CO_2_] (*C*_i_) under saturating light (1500 μmol photons m^−2^ s^−1^). Values show the mean ± SEM of the three separate EP1S1S2 lines, each comprised of 5-8 individual measurements on separate rosettes (see **Fig. S10** and **S11** for further details). **D.** Violin plot showing the proportion of starch area associated with the condensate (enclosed or adjacent), in EP1S1S2-1 (Line 1), representative of 45-48 starch-containing chloroplasts. Asterisks show significance where P < 0.001 using a Mann Whitney test. **E.** Representative TEM images from the pre-dawn dataset analysed in **D**. Scale bar = 0.5 µm.

We hypothesised that the large condensate-associated starch granules in the EP1S1S2 lines could be metabolised more slowly, which might account for their reduced growth phenotype. To determine this, an additional pre-dawn TEM analysis was carried out on line EP1S1S2-1. Here we observed a greater proportion of condensate-associated starch remained in the pre-dawn assay than at the end of the day (**Fig. 5*D*** and **5*E***). Thus, the growth phenotype observed in the EP1S1S2 lines could be a result of inefficient starch remobilisation during the dark period, likely due to the size and shape of the large, rounded condensate-associated granules, which would have a reduced surface area relative to lenticular stromal starch granules.

## Discussion

In this study we have taken a crucial step forward in the development of a functional pCCM to enhance the efficiency of photosynthesis in a C3 plant. A starch sheath diffusion barrier is predicted to improve the efficiency of the Chlamydomonas pCCM by increasing the concentration of CO_2_ around Rubisco by up to 10-fold (14). This is supported by observations of starch sheath mutants in Chlamydomonas, which show reduced growth in low CO_2_ conditions (13). SAGA1 and SAGA2 localise to the pyrenoid and form an interface between the Rubisco matrix and the starch sheath via RBMs and putative CBM20 domains, respectively (21–23). Based on previous work, SAGA1 was an ideal candidate to recruit starch to the proto-pyrenoid condensate in Arabidopsis (21), with the aim of generating a starch sheath to limit CO_2_ diffusion as part an efficient functional pCCM (15). SAGA2 has remained largely uncharacterised to date, but previous work has suggested it could augment the formation of starch around the proto-pyrenoid (22). We found that SAGA1 did indeed recruit starch to the proto-pyrenoid condensate in Arabidopsis, including extended plate-like starch granules at the edge of the condensate. But SAGA1 alone was not sufficient for the recruitment of multiple adjacent starch granules. Subsequently, we showed that co-expression of SAGA2 led to a further significant increase in condensate-associated starch including the formation of a ring of adjacent starch granules.

The different localisation patterns observed for SAGA1 and SAGA2 when expressed heterologously *in planta* suggested that they have distinct roles (22). SAGA1 is critical for the formation of a single pyrenoid condensate and a proper starch sheath in Chlamydomonas (21). Recent work has suggested that SAGA1 also contributes to the appropriate localisation of pyrenoid-associated CCM components, such as LCIB and the Ca^2+^-binding protein CAS, which in turn impacts CAS-dependent retrograde regulation of several nuclear encoded pCCM components (44). Notably, we observed that SAGA1 expression resulted in the formation of a scaffold-like structure within the proto-pyrenoid condensate. In contrast, SAGA2 did not produce any patterns in the condensate and appeared to preferentially bind starch, which resulted in localisation to non-condensate starch or even splitting of the condensate likely through binding to multiple starch granules. Our data suggest that the primary role of SAGA2 is as a starch-recruiting scaffold between the Rubisco matrix and starch sheath in Chlamydomonas. This hypothesis is supported by the differing localisation patterns of SAGA1 and SAGA2 in Chlamydomonas; SAGA1 has a punctate localisation around the pyrenoid periphery, whilst the homogeneous expression pattern along the periphery for SAGA2 suggests a more prominent role in the matrix-starch interface (22). We confirmed that SAGA2 also interacts with EPYC1 through the same alpha-helices that facilitate interaction between EPYC1 and the small subunit of Rubisco (19, 26). The functional significance of being able to interact both with Rubisco and EPYC1 is unclear, but this may help to anchor SAGA2 to the pyrenoid if SAGA2 has a high starch-binding affinity.

Starch granules associated with the condensate (enclosed or adjacent) were morphologically distinct from stromal starch granules. Starch granule initiation and growth in Arabidopsis mesophyll chloroplasts occurs in stromal pockets between the thylakoids (29), thus the morphology of granules can be linked to physical factors, such as correct thylakoid organisation (33). In addition to physical factors, biochemical factors like STARCH SYNTHASE 4 (SS4) not only influences the size and number of starch granules, but also regulate the anisotropic growth of granules, leading to their lenticular shape (29, 45). SS4, along with other components that are important for establishing correct granule number, such as MAR1-BINDING FILAMENT PROTEIN 1 (MFP1) and PROTEIN TARGETING TO STARCH 2 (PTST2), are at least partially associated with thylakoid membranes (32, 46). We hypothesise that SAGA1 recruited newly initiated starch granules to the proto-pyrenoid condensate. When enclosed within the condensate, the developing granule can grow, presumably due to the ability of starch synthesis proteins to diffuse into the matrix. However, the lack of contact with the thylakoids may allow free isotropic growth without the physical restrictions imposed by the membranes, or could prevent access to SS4 and other thylakoid-bound proteins that are important for granule morphogenesis. Similarly, the curvature of adjacent granules on the surface of the condensate is likely due to a lack of thylakoid contact and physical restrictions imposed by the condensate. SAGA2 led to the further recruitment of granules to the condensate, which grew into the large and irregularly shaped granules observed in EP1S1S2 lines. Future work should focus on clarifying the sites of initiation for condensate-associated starch granules, the contribution of SS4 and other starch synthesis proteins to their shape, as well as identify the proteins involved in their breakdown.

The plate-like starch granules surrounding the proto-pyrenoid in EP1S1S2 lines could provide the foundations of a diffusion barrier for an efficient functional pCCM in plants. To improve coverage over the surface of the condensate, additional proteins will need to be employed to help remodel the starch and provide a more complete barrier, and allow more efficient remobilisation of starch so as to not negatively impact on growth. Although fluorescent proteins by themselves do not appear to affect starch metabolism or granule morphology (47), the presence of fluorescent tags on EPYC1, SAGA1 or SAGA2 in our current plant lines may restrict capacity for screening additional components, or the use of fluorophores such as fluorescein for more high-throughput analyses of leaf starch localisation and structure (48). Thus, future work will focus on generating fully untagged EP1S1S2 lines to screen proteins of potential interest, including starch re-modelling proteins such as isoamylases or glucanotransferases (13), starch branching enzymes SBE3 and SBE4, which surrounds the pyrenoid and interact with Rubisco, respectively, or pyrenoid-peripheral proteins such as LCI9, which is proposed to form a mesh between the starch plates (18).

## Materials and Methods

### Plant material and growth conditions

Arabidopsis (*Arabidopsis thaliana*, Col-0 background) seeds were sown on compost, stratified for 3 d at 4 °C and grown at 20 °C, ambient CO_2_ and 70% relative humidity under 200 μmol photons m^-2^ s^-1^ supplied by cool white LED lights (Percival SE-41AR3cLED, CLF PlantClimatics GmbH, Wertingen, Germany) in 12 h light and 12 h dark, or 16 h light and 8 h dark. For comparisons of different genotypes, plants were grown from seeds of the same age and storage history, harvested from plants grown in the same environmental conditions.

### Plasmid vector design and transformation

The coding sequence of SAGA1 and SAGA2 were codon optimised for expression in Arabidopsis using GeneArt software (Thermo Fisher Scientific), and cloned directly into the level 0 acceptor vectors pAGM1287 and pAGM1299, respectively, of the Plant MoClo system (49). To generate fusion proteins, gene expression constructs were assembled into binary level 2 acceptor vectors (see **Table S1** for further vector details and **Fig. S12** for vector maps). The 35S cauliflower mosaic virus (CaMV) promoter and CsVMV (cassava vein mosaic virus) promoter were used to drive expression. Level 2 vectors were transformed into *Agrobacterium tumefaciens* (AGL1) for stable insertion in Arabidopsis plants by floral dipping (50). Homozygous transgenic and azygous lines were identified in the T2 generation using the pFAST-R selection cassette or kanamycin resistance (51).

### Protein analyses

Soluble protein was extracted from frozen leaf material of 21-d-old plants (sixth and seventh leaf) in protein extraction buffer (20 mM Tris-HCl pH 7.5 with 5 mM MgCl_2_, 300 mM NaCl, 5 mM DTT, 1% Triton X-100 and cOmplete Mini EDTA-free Protease Inhibitor Cocktail (Roche, Basel, Switzerland)). Samples were heated at 80 °C for 15 min with 1 x Bolt LDS sample buffer (ThermoFisher Scientific, UK) and 200 mM DTT. Extracts were centrifuged and the supernatants subjected to SDS-PAGE on a 12% (w/v) polyacrylamide gel and transferred to a nitrocellulose membrane. Membranes were probed with rabbit serum raised against wheat Rubisco at 1:10,000 dilution (52), the SSU RbcS2 from Chlamydomonas (CrRbcS2) (raised to the C-terminal region of the SSU (KSARDWQPANKRSV) by Eurogentec) at 1:1,000 dilution, ACTIN (AS21 4615, Agrisera) at 1:1000 dilution, SAGA1 at 1:1000 dilution (21), SAGA2 (raised against the peptide sequence SRNGTHAAGEDVREV by Eurogentec) at 1:1000 and EPYC1 at 1:2,000 dilution (18), followed by HRP-linked goat anti-rabbit IgG (Abcam) or HRP-linked rabbit anti-mouse (Agrisera) at 1:10,000 dilution, and visualised using Pierce ECL Western Blotting Substrate (Life Technologies).

### Condensate extraction

Soluble protein was extracted as before, then filtered through Miracloth (Merck Millipore, Burlington, Massachusetts, USA), and centrifuged at 500 *g* for 3 min at 4 °C, as in Mackinder et al. (18). The pellet was discarded, and the extract was centrifuged again for 12 min. The resulting pellet was washed once in protein extraction buffer, then re-suspended in a small volume of buffer and centrifuged again for 5 min. Finally, the pellet was re-suspended in 25 µl of extraction buffer and used in confocal analysis or SDS-PAGE electrophoresis.

### Growth analysis and photosynthetic measurements

Rosette growth rates were quantified using an in-house imaging system (53). Maximum quantum yield of photosystem II (PSII) (*F*_v_/*F*_m_) was measured on 32-d-old plants using a Hansatech Handy PEA continuous excitation chlorophyll fluorimeter (Hansatech Instruments Ltd, King’s Lynn, UK) (54). Gas exchange and chlorophyll fluorescence were determined using a LI-COR LI-6400 (LI-COR, Lincoln, Nebraska, USA) portable infra-red gas analyser with a 6400-40 leaf chamber on either the sixth or seventh leaf of 35- to 45-d-old non-flowering rosettes grown in large pots under 200 μmol photons m^-2^ s^-1^ to generate leaf area sufficient for gas exchange measurements (55). The response of *A* to the intercellular CO_2_ concentration (*C*_i_) was measured at various CO_2_ concentrations (50, 100, 150, 200, 250, 300, 350, 400, 600, 800, 1000 and 1200 µmol mol^−1^) under saturating light (1,500 μmol photons m^-2^ s^-1^) and ambient O_2_ concentrations or 2% O_2_. For all gas exchange experiments, the flow rate was kept at 200 μmol mol^-1^, leaf temperature was controlled at 25 °C and chamber relative humidity was maintained at *ca*. 70%. Measurements were performed after net assimilation and stomatal conductance had reached steady state. Gas exchange data were corrected for CO_2_ diffusion from the measuring chamber as in Bellasio et al. (56). To estimate *V*_cmax_, *J*_max_, the CO_2_ compensation point (*Γ*) and mesophyll conductance to CO_2_ (*g*_m_) the *A*/*C*i data were fitted to the C_3_ photosynthesis model as in Ethier and Livingston (57) using the catalytic parameters *K*_c_^air^ and affinity for O_2_ (*K*_o_) values for wild-type Arabidopsis Rubisco at 25 °C and the Rubisco content of WT and S2_Cr_ lines (36).

### Confocal laser scanning and fluorescence recovery after photobleaching

Leaves were imaged with a Leica TCS SP8 laser scanning confocal microscope (Leica Microsystems, Milton Keynes, UK) as in Atkinson et al. (35). Processing of images was done with Leica LAS AF Lite software. Fluorescence recovery after photobleaching (FRAP) was carried out with a 40x water immersion objective and a photomultiplier tube (PMT) detector. The 488 nm laser was set to 2% power for pre- and post-bleach images, and 80% for the bleaching step. Pre- and post-bleach images were captured at 328 ms intervals. For photo-bleaching the laser was directed to a region with a diameter of 0.5-0.6 µm on one side of the EPYC1 condensate. Recovery time was calculated by comparing GFP expression to an unbleached region of the same size.

### Immunogold labelling, PATAg staining and transmission electron microscopy

Leaf samples were taken from 21-d-old plants and fixed with 4% (v/v) paraformaldehyde, 0.5% (v/v) glutaraldehyde and 0.05 M sodium cacodylate (pH 7.2). Leaf strips (1 mm wide) were vacuum infiltrated with fixative three times for 15 min, then rotated overnight at 4 °C. Samples were rinsed three times with PBS (pH 7.4) then dehydrated sequentially by vacuum infiltrating with 50%, 70%, 80% and 90% ethanol (v/v) for 1 hr each, then three times with 100%. Samples were infiltrated with increasing concentrations of LR White Resin (30%, 50%, 70% (w/v) in ethanol) for 1 hr each, then 100% resin three times. The resin was polymerised in capsules at 50 °C overnight. Sections (1 μm thick) were cut on a Leica Ultracut ultramicrotome, stained with Toluidine Blue, and viewed in a light microscope to select suitable areas for investigation. Ultrathin sections (60 nm thick) were cut from selected areas and mounted onto plastic-coated copper grids. Grids were stained in 2% (w/v) uranyl acetate then viewed in a JEOL JEM-1400 Plus TEM (JEOL, Peabody, Massachusetts, USA). Images were collected on a GATAN OneView camera (GATAN, Pleasanton, California, USA). For immunogold labelling, grids were blocked with 1% (w/v) BSA in TBSTT (Tris-buffered saline with 0.05% (v/v) Triton X-100 and 0.05% (v/v) Tween 20), incubated overnight with anti-SAGA1 antibody in TBSTT at 1:10 dilution, and washed twice each with TBSTT and water. Incubation with 15 nm gold particle-conjugated goat anti-rabbit secondary antibody (Abcam, Cambridge, UK) in TBSTT was carried out for 1 hr at 1:10 dilution, before washing as before. Starch was visualised by TEM using the periodic acid-thiocarbohydrazide-silver proteinate (PATAg) staining method. Sections were mounted on grids and floated on 1% (v/v) periodic acid in water for 30 mins, washed in water three times for 15 mins, then floated on 0.2% (v/v) thiosemicarbazide in 20% (v/v) acetic acid for 24 h. Grids were washed as before, then floated on 1% (w/v) silver proteinate in water for 30 mins, then washed again before staining in 2% (w/v) uranyl acetate and visualising.

### SBF-SEM sample preparation and image acquisition

Samples were fixed overnight in 2.5% (v/v) glutaraldehyde (GA), 2% (v/v) paraformaldehyde (PFA) and 2 mM CaCl_2_ in 0.1 M sodium cacodylate buffer (NaCaC), pH 7.4. Fixed samples were then placed in fresh NaCac buffer inside a Pelco BioWave Pro+ (Agar Scientific) for subsequent irradiation under vacuum during the staining protocol (see steps 1 and 11 in **Table S2**). Durcupan ACM resin mixture was prepared according to manufacturer’s instructions (Sigma-Aldrich) with components A (epoxy resin), B (hardener), C (accelerator) and D (dibutyl phthalate). After step 11 of the infiltration, the samples were removed from the BioWave and placed in fresh 100% Durcupan resin (Merk Life Sciences) and left overnight on a rotator in the fume hood (step 12). The following day, samples were placed into 8 mm flat polypropylene TAAB capsules containing fresh Durcupan resin and polymerised in the BioWave Pro+ for 30 min at 60°C, then 90 min at 100 °C (steps 13 and 14).

Samples were manually trimmed to size using a razor blade then adhered to a Gatan 3View specimen pin, standard type 1.4 mm flat (Agar Scientific Ltd), using conductive epoxy glue. A Leica ARTOS 3D Ultramicrotome (Leica microsystems) was used with a DiATOME trim 45 diamond trimming tool (Leica microsystems) to refine the geometry of the resin block. Sample pins were subsequently sputter coated with 15 nm of platinum using a Leica ACE 600 sputter coater (Leica microsystems). The samples were then placed in a Zeiss Gemini 300 SEM (Carl Zeiss Microscopy) with focal charge compensation, equipped with a Gatan 3View® 2XP in-situ ultramicrotome and OnPoint detector (AMETEK).

Images were captured at 1.5 kV under high vacuum conditions, using Gatan DigitalMicrograph DMS3 software (AMETEK). Sections thickness was 50 nm, pixel size ranged from 25-50 nm, pixel dwell time ranged from 10-20 µs and image size 1040 x 1042 or 2048 x 2048. Datasets were aligned and segmented on Microscopy Image Browser (MIB) (http://mib.helsinki.fi), and final images were visualised using Amira software (Thermo Fisher Scientific).

### Starch area measurements and content assays

Starch area analysis was carried out by measurement of TEM images using Fiji (ImageJ). Chloroplast and starch granule area was derived using the freehand selection tool and the area function. Circularity was assessed using the circularity function, which uses the formula: *circularity = 4π (area / perimeter^2^).* Enzymatic starch content assays were carried out on 26-d-old plants grown in 16 h light : 8 hr dark cycles. Harvested rosettes were ground in liquid nitrogen and boiled in 70 % (v/v) ethanol for 10 min at 90 °C, before centrifuging at 5000 *g* for 10 min at 4 °C. The supernatant was removed and the pellet re-suspended in sterile water, before centrifuging again and discarding the supernatant. The pellet was re-suspended in 1.5 ml of 80% (v/v) ethanol and boiled for 10 min at 90 °C, then centrifuged and the supernatant removed. The pellet was washed again in 80% (v/v) ethanol and allowed to air-dry, before re-suspending in sterile water to 1 ml. The sample was boiled at 95 °C for 20 min to gelatinise the starch, then 200 µl of each sample was digested for 16 h at 37 °C with α-amyloglucosidase (6 U/ml, Merck), and α-amylase (4 U/ml, Merck) in 200 mM sodium acetate– acetic acid, pH 4.8, before centrifuging at 14,000 *g* for 10 min at 20 °C. To determine the glucose concentration, 5 µl sample was mixed with 245 µl assay cocktail (122 mM Tris, pH 8.1, 4.1 mM MgCl_2_, 50 µg NAD, 165 µg ATP), 0.3 U hexokinase (Scientific Laboratory Supplies) and 1 U glucose-6-phosphase dehydrogenase (Merck) in triplicate. The absorbance at 340 nm was recorded before and after the reaction, and the concentration of glucose calculated by comparing with a standard curve.

### Yeast two-hybrid

Yeast two-hybrid was used to detect interactions between proteins of interest as described previously (26). Briefly, genes were cloned into pGBKT7 and pGADT7 vectors to create fusions with the GAL4-DNA binding domain or activation domain, respectively (see **Table S3** for gene sequences). Competent yeast cells (Y2H Gold, Clontech) were co-transformed with the two vectors using a polyethylene glycol (PEG) solution (100 mM LiAc, 10 mM Tris–HCl pH 7.5, 40% (v/v) PEG 4000) with 167 µg/ml DNA from salmon sperm. Cells were incubated at 30 °C for 30 min, then subjected to a heat shock of 42 °C for 20 min. After 90 mins of recovery in YPDA, cells were plated onto SD-L-W (standard dextrose medium lacking leucine and tryptophan; Anachem). Ten to fifteen of the resulting colonies were pooled, cultured, and plated onto SD-L-W and SD-L-W-H (Anachem) containing different concentrations of the HIS3 inhibitor tri-aminotriazole (3-AT), and incubated for 3 d before assessing for the presence or absence of growth.

### Statistical analyses

Results were subjected to one-way analysis of variance (ANOVA) to determine the significance of the difference between sample groups. When one-way ANOVA was performed, Tukey’s honestly significant difference (HSD) post-hoc tests were conducted to determine the differences between the individual treatments (IBM SPSS Statistics, Ver. 26.0).

## Acknowledgments

This work was funded by the UK Research and Innovation Biotechnology and Biological Sciences Research Council (BB/S015531/1 and BB/W003538/1) and Leverhulme Trust (RPG-2017-402). TEM was carried out with the support of Stephen Mitchell and the Wellcome Trust Multi User Equipment Grant (WT104915MA). SBF-SEM work was funded through BBSRC (BB/X001520/1) and the BBSRC-funded Institute Strategic Programme Harnessing Biosynthesis for Sustainable Food and Health (HBio)(BB/X01097X/1). We thank JIC Bioimaging for providing access to microscopes.

## Supplementary Figures and Tables

**Figure S1.**
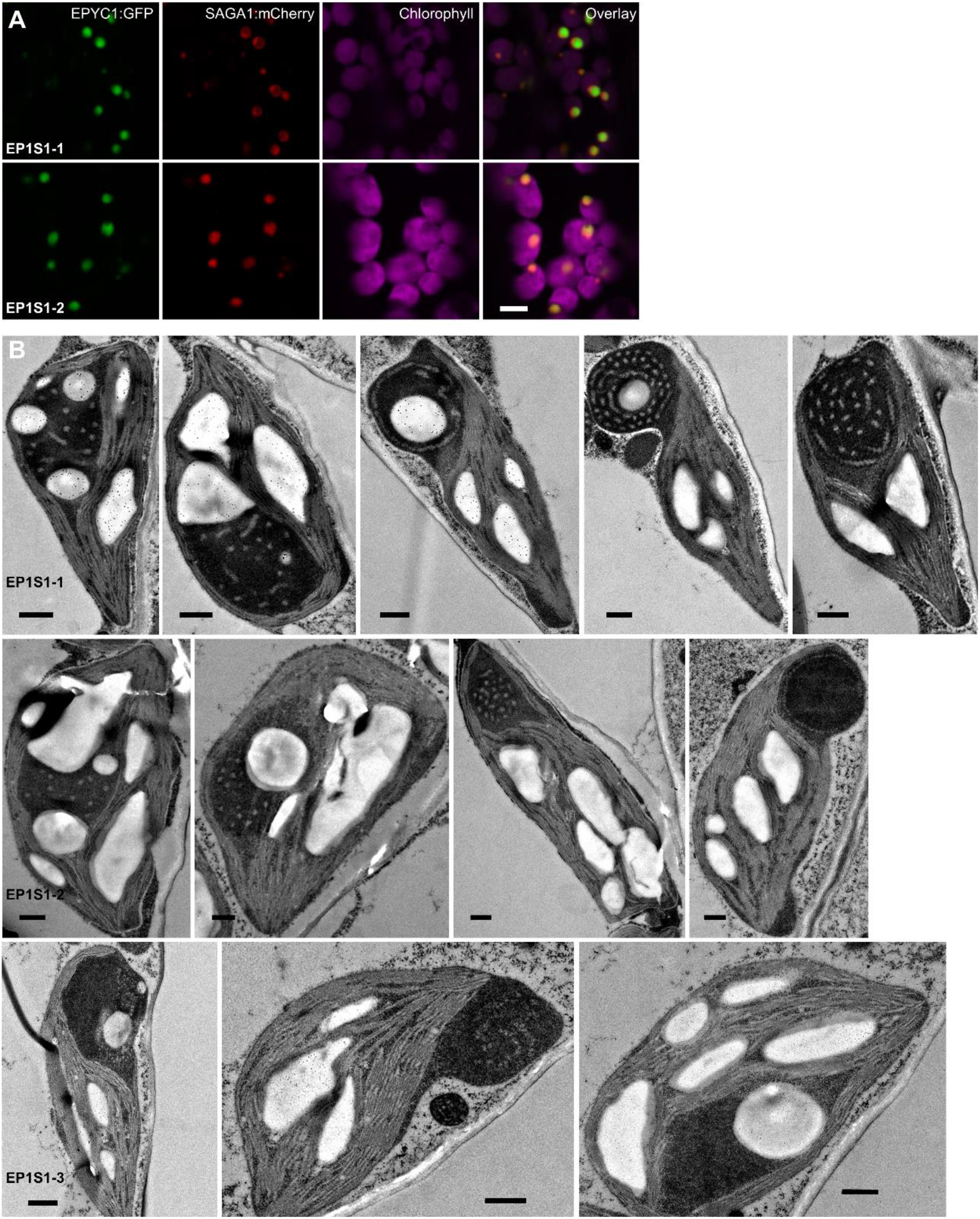
Additional confocal and TEM images for EP1S1 lines. **A.** Confocal image examples of alternative expression patterns for SAGA1::mCherry localised to the proto-pyrenoid from lines EP1S1-1 and EP1S1-2 (Fig. 1D). Scale bar = 0.5 µm. **B.** Representative TEM image examples for EP1S1-1, EP1S1-2 and EP1S1-3. Scale bar = 0.5 µm.

**Figure S2.**
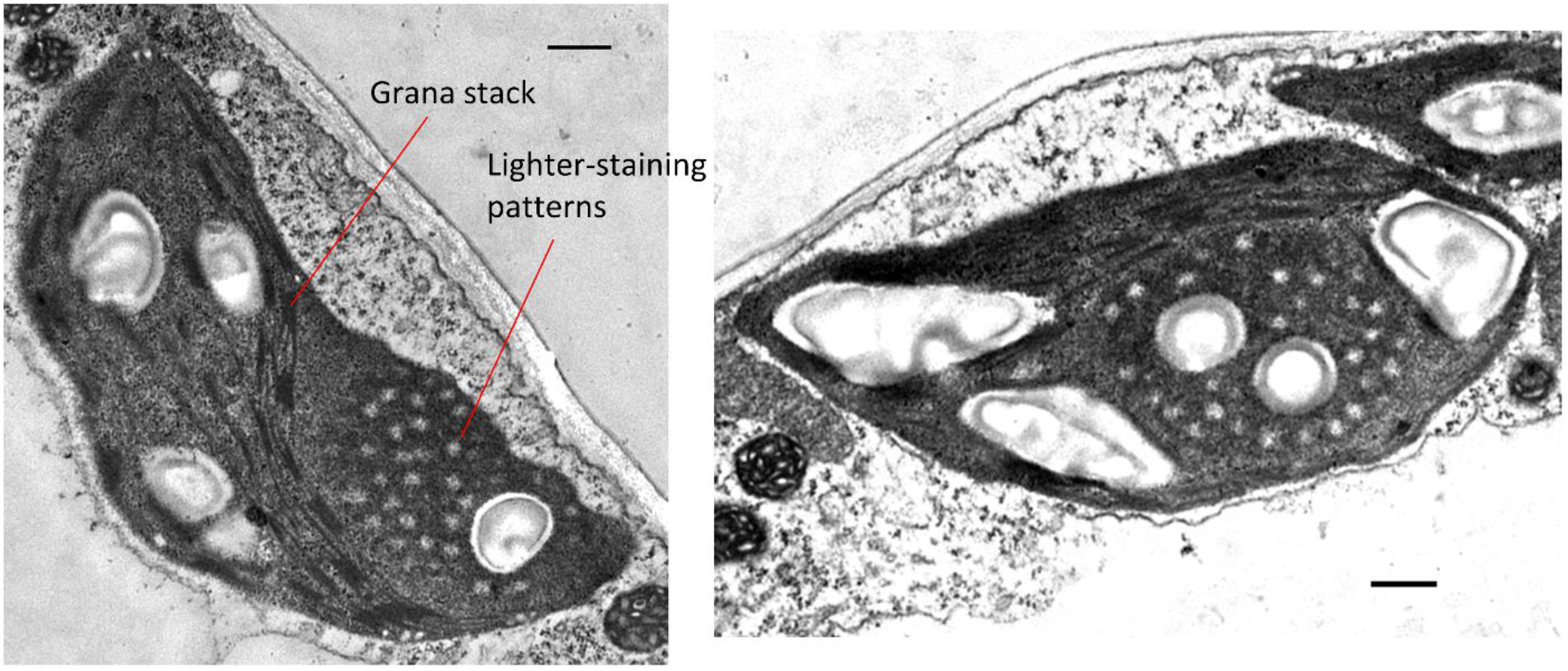
Lighter-staining regions in EP1S1-1 are not stained by osmication. Following fixation EP1S1-1 leaf samples were stained with 1% (w/v) osmium tetroxide in 0.05 M sodium cacodylate for 45 minutes. Samples were washed three times with phosphate buffered saline, before being dehydrated and embedded as normal. Membranes such as the visible grana stacks were stained black during this process, whilst the patterned structures in the condensate matrix remained lighter-coloured. Scale bar = 0.5 µm.

**Figure S3.**
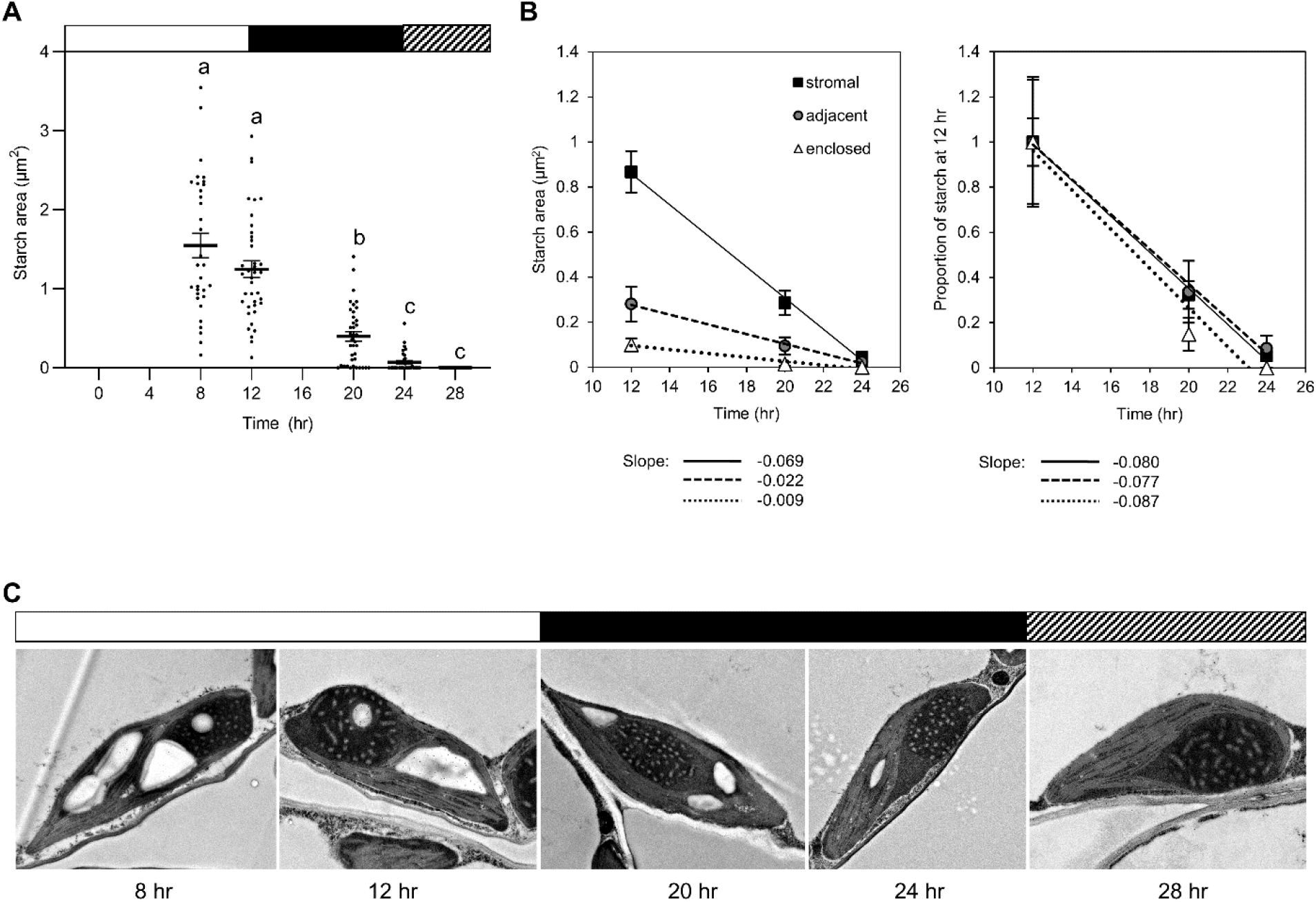
Total starch area and representative images in EP1S1-1 over time. **A.** Average of total areas for stroma, adjacent and enclosed starch granules in EP1S1-1 plants over time from Fig. 2G. The averages at each time point represents the mean ± SEM of 22-39 images. Letters indicate significant difference (p < 0.05) as determined by one-way ANOVA followed by a Kruskal-Wallis post-hoc test. **B.** Starch degradation rates plotted for each starch type showing total starch area (left) and proportional starch area (right). Trendlines are fitted to the data and the values for the slopes given under each graph. **C.** Representative TEM images of starch abundance over time.

**Figure S4.**
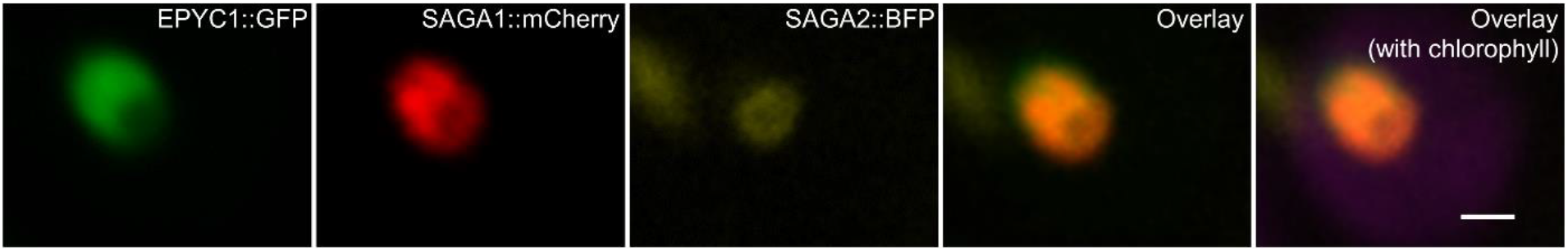
Representative confocal images of plants expressing EPYC1, SAGA1 and SAGA2 (EP1S1S2) with each transgene tagged with a unique fluorophore. Scale bar = 1 µm.

**Figure S5.**
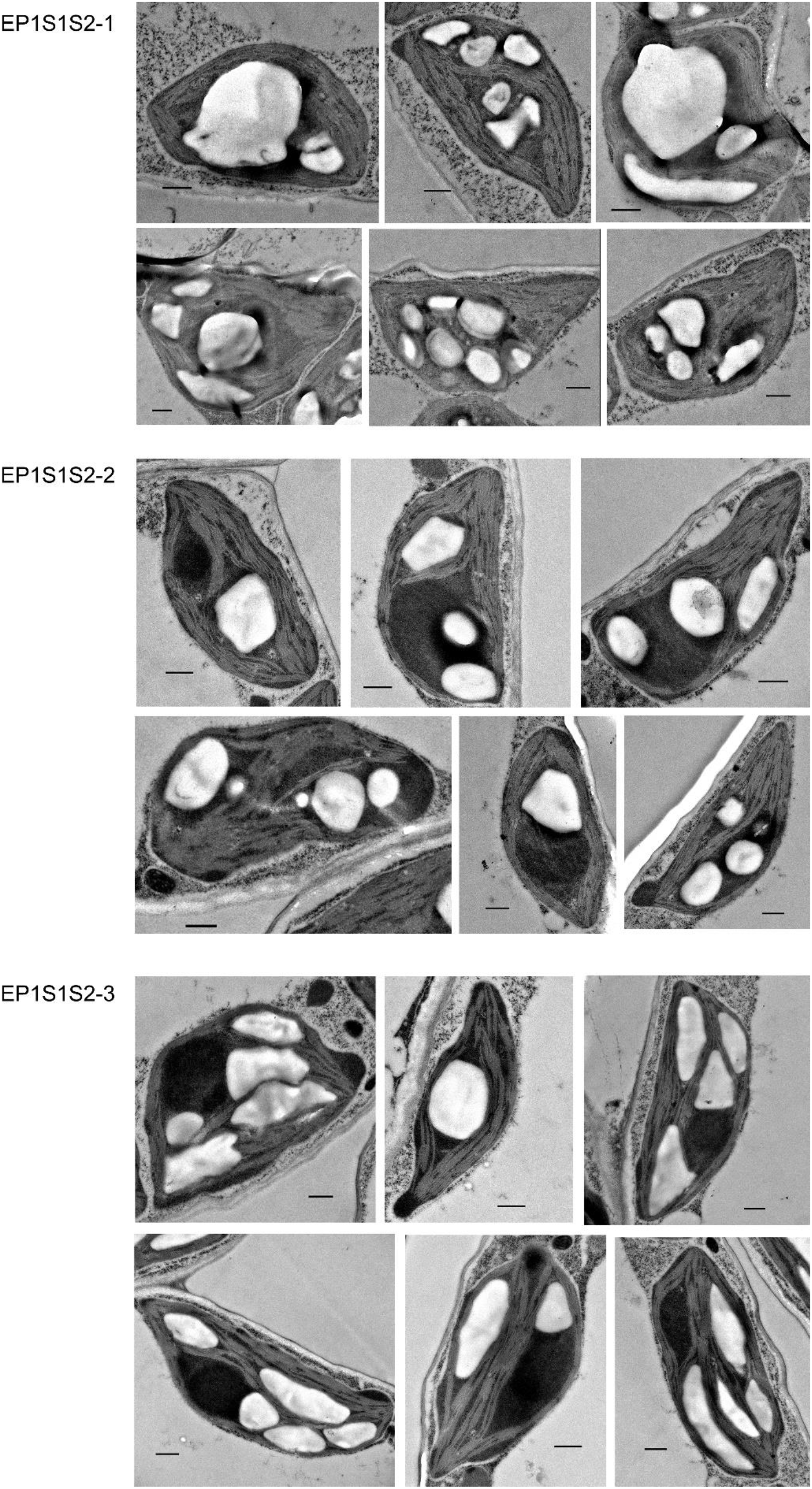
Additional representative TEM images for each of the three EP1S1S2 lines. See Fig. 3 for transgene expression levels. Scale bar = 0.5 µm.

**Figure S6.**
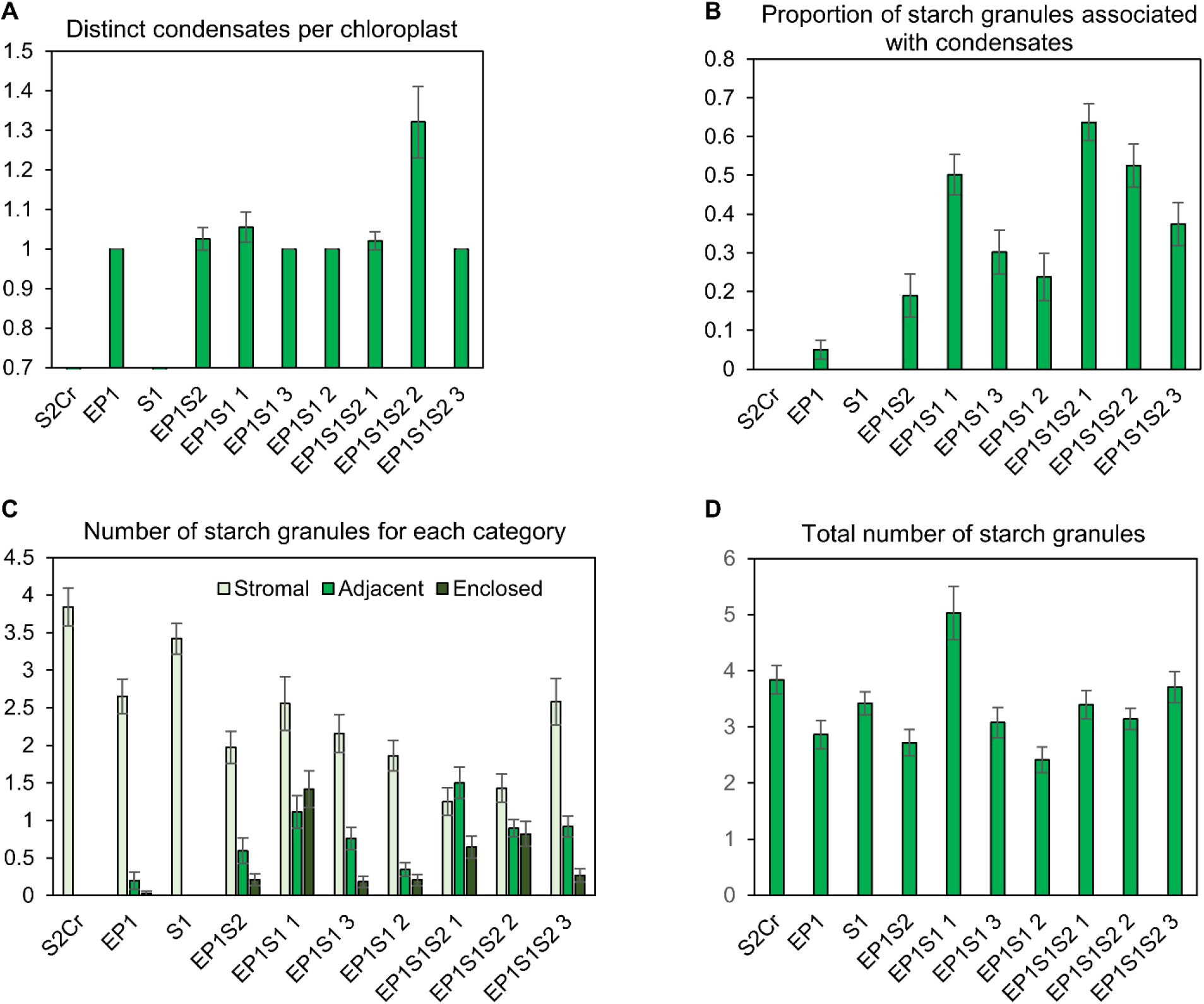
Additional starch granule parameters from area analysis of TEM images. **A.** The number of distinct condensates observed per chloroplast in each line. **B.** Proportion of total starch granules per chloroplast that are associated with the condensate (adjacent or enclosed granules) compared to those that were located in the stroma (stromal granules). **C.** Total number of starch granules per chloroplast classified as stromal, adjacent or enclosed. **D.** Total number of starch granules per chloroplast. Error bars show the mean ± SEM of 24-48 chloroplasts per line.

**Figure S7.**
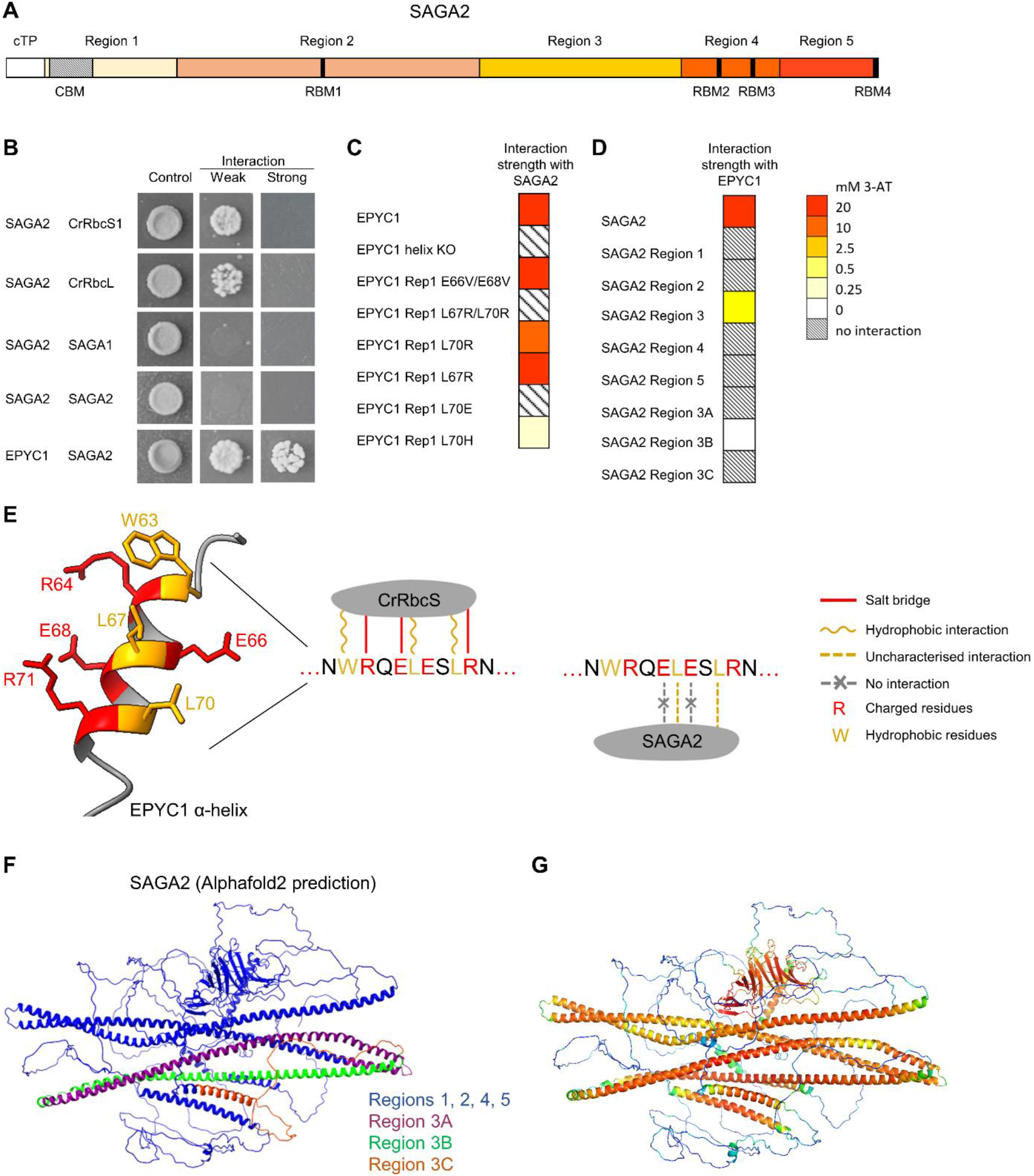
SAGA2 interacts with EPYC1 through hydrophobic residues on the EPYC1 α-helix. **A.** SAGA2 (Cre09.g394621, 1817 amino acid residues) contains a C-terminal carbohydrate binding motif (CBM) and four Rubisco binding motifs (RBMs, black bars). A predicted chloroplast transist peptide (cTP) and regions for interaction analysis (below) are shown. **B.** Yeast 2-hybrid (Y2H) assays supported protein-protein interactions between SAGA2 and the small subunit 1 of Rubisco in Chlamydomonas (CrRbcS1; Cre02.g120100), the large subunits of Rubisco (CrRbcL) and EPYC1. Control column shows growth on CSM-L-W, whilst interaction columns show growth on CSM-L-W-H dropout media (weak), and with 10 mM 3-aminotriazole (3-AT) inhibitor (strong). **C.** Y2H assays showing the interaction strength between SAGA2 and EPYC1 with mutations in various residues of the α-helix. Interaction strength was scored by growth on increasing concentrations of 3-AT as in Atkinson et al. (26). The EPYC1 α-helix KO is a full length EPYC1 in which each of five α-helix sequences was replaced by alanine residues. In the subsequent samples just one repeat of EPYC1 is tested for interaction (Rep1, corresponding to amino acid residues 28–76 of the full-length protein), with one or more residues substituted. **D.** SAGA2 was split into 5 regions, with region 3 further subdivided into three parts (A-C), to test for interaction with EPYC1. **E.** Model of one of the repeat α-helix sequences of EPYC1 (residues 63-72) with hydrophobic residues shown in orange and charged residues shown in red. A comparison of the residues that interact with CrRbcS (19) and SAGA2 (from data in C) is shown. **F.** Predicted structure of SAGA2 (Alphafold2), with sections of region 3 highlighted in magenta (A), green (B) and orange (C), respectively. **G**. Predicted SAGA2 structure coloured by confidence score according to the Predicted Local Difference Distance Test (pLDDT), where red = high and blue = low. Please note that SAGA2 is predicted to contain several intrinsically disordered regions (IDRs), and the structural predictions of AlphaFold can be unreliable for IDRs.

**Figure S8. SBF-SEM raw video data and 3-D reconstruction videos** (*attached with submission*). **A, B.** Raw and 3-D reconstruction videos for the SAGA1 network in EP1S1 in Fig. 4E (‘Video - raw for Fig 4E.mp4’ and ‘Video - 3-D reconstruction for 4E.mp4’). **C, D.** Videos for stromal, enclosed and adjacent starch granules in EP1S1 in Fig. 4F (‘Video - raw for Fig 4F.mp4’ and ‘Video - 3-D reconstruction for 4F.mpg’). **E, F.** Videos for adjacent starch granules in EP1S1S2 in Fig. 4G (‘Video - raw for Fig 4G.mp4’ and ‘Video - 3-D reconstruction for 4G.mpg’). **G, H.** Videos for EP1-1 in Fig. 1C (‘Video - raw for EP1.mp4’ and ‘Video - 3-D reconstruction for EP1.mp4’). **I, J.** Videos for S1-1 in Fig. 1C (‘Video - raw for S1-1.mp4’ and ‘Video - 3-D reconstruction for S1-1.mp4’). **K, L.** Videos for S2_Cr_ in Fig. 1C (‘Video - raw for S2Cr.mp4’ and ‘Video - 3-D reconstruction for S2Cr.mp4’).

**Figure S9.**
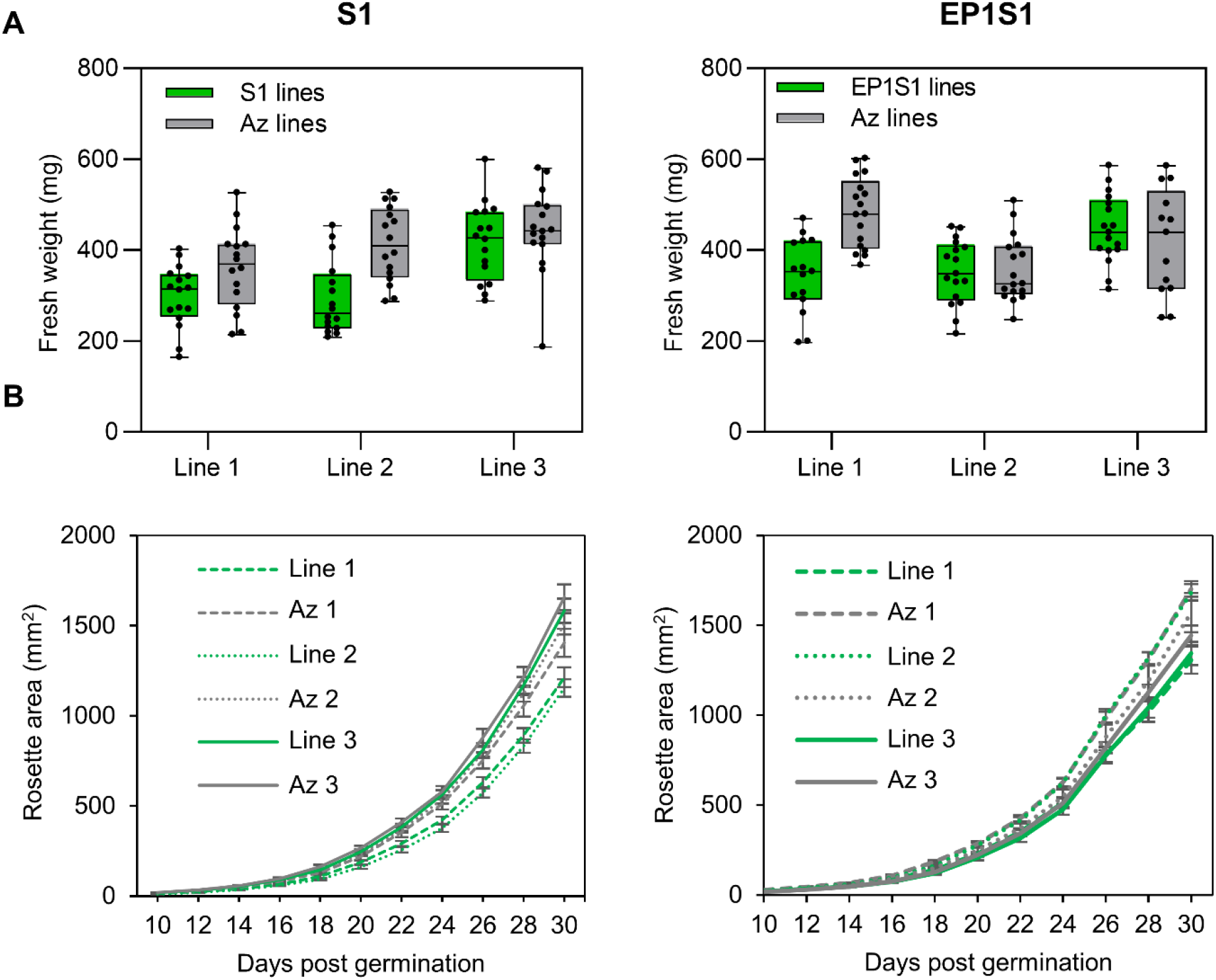
Fresh weight and rosette areas of S1 and EP1S1 lines. **A.** Fresh weight of S1 and EP1S1 lines and their azygous (Az) segregants 32 days after germination. **B.** Rosette expansion for S1 and EP1S1 lines measured over 30 days post germination. Error bars show the mean ± SEM of 12-21 individual rosettes.

**Figure S10.**
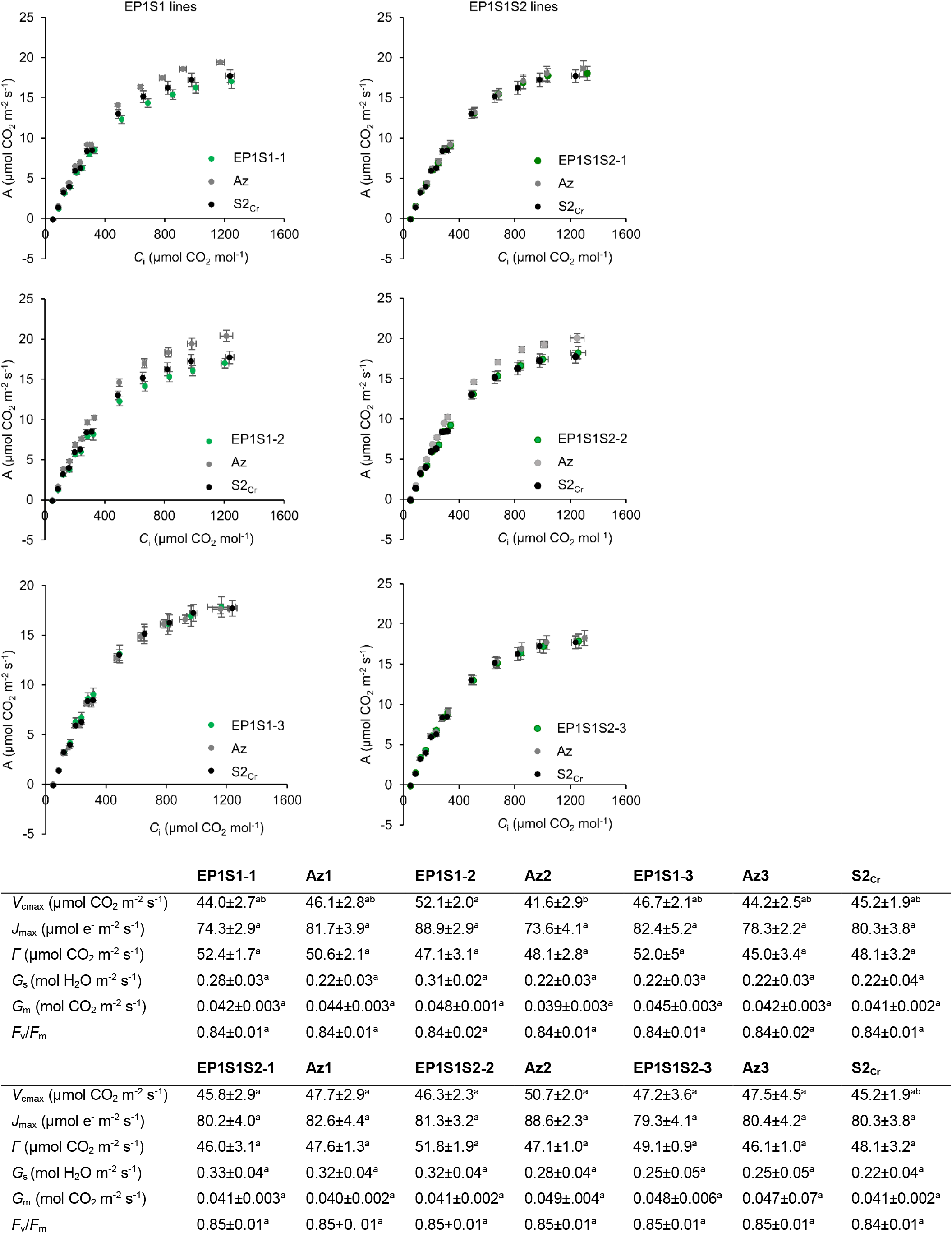
Net CO_2_ assimilation (*A*) based on sub-stomatal [CO_2_] (*C*_i_) under saturating light (1500 μmol photons m^−2^ s^−1^) for EP1S1 and EPS1S2 lines. Values show the mean ± SEM of 5-8 individual measurements on separate rosettes for EP1S1 (left) and EP1S1S2 (right) plants, their azygous segregants and the background line S2_Cr_. Variables derived from gas exchange data include maximum rate of Rubisco carboxylation (*V*_cmax_), maximum electron transport rate (*J*_max_), CO_2_ compensation point (*Γ*) , stomatal conductance (*G*_s_), mesophyll conductance (*G*_m_) and the maximum quantum yield of photosystem II (*F*_v_/*F*_m_). Letters indicate significant difference (p < 0.05) as determined by one-way ANOVA followed by Tukey’s honestly significant difference (HSD) post-hoc tests.

**Figure S11.**
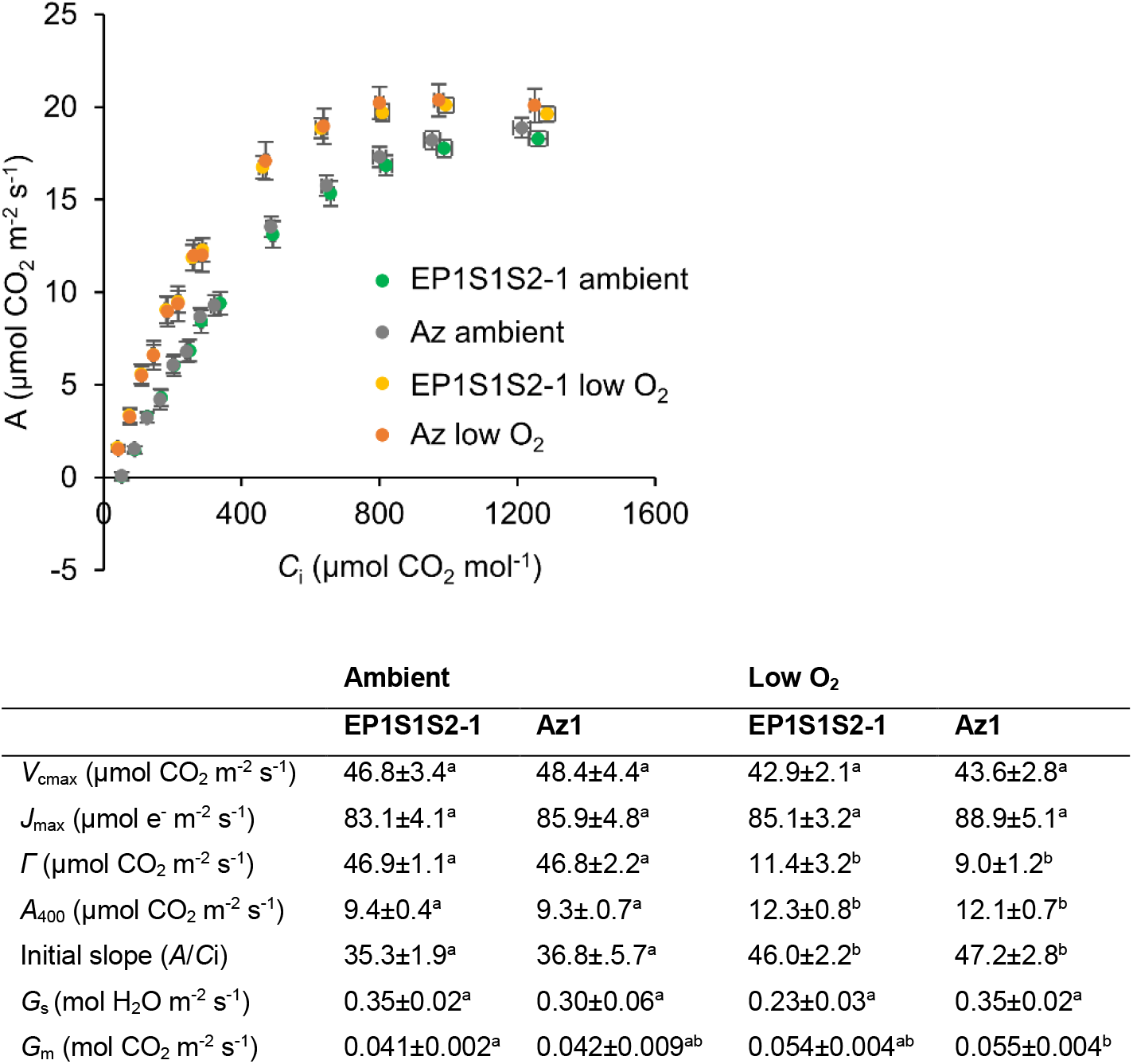
Net CO_2_ assimilation (*A*) based on sub-stomatal [CO_2_] (*C*_i_) under saturating light (1500 μmol photons m^−2^ s^−1^) for EPS1S2 under photorespiratory and non-photorespiratory conditions. Values show the mean ± SEM of 6 individual measurements on separate rosettes for EP1S1S2-1 plants and azygous segregants measured under ambient O_2_ and 2% O_2_. Variables derived from gas exchange data include maximum rate of Rubisco carboxylation (*V*_cmax_), maximum electron transport rate (*J*_max_), CO_2_ compensation point (*Γ*), CO_2_ assimilation rate at near ambient CO_2_ (*A*_400_), carboxylation efficiency taken as the initial slope between 50 and 350 µmol mol^−1^ ambient CO_2_, stomatal conductance (*G*_s_) and mesophyll conductance (*G*_m_). Letters indicate significant difference (p < 0.05) as determined by one-way ANOVA followed by Tukey’s honestly significant difference (HSD) post-hoc tests.

**Figure S12.**
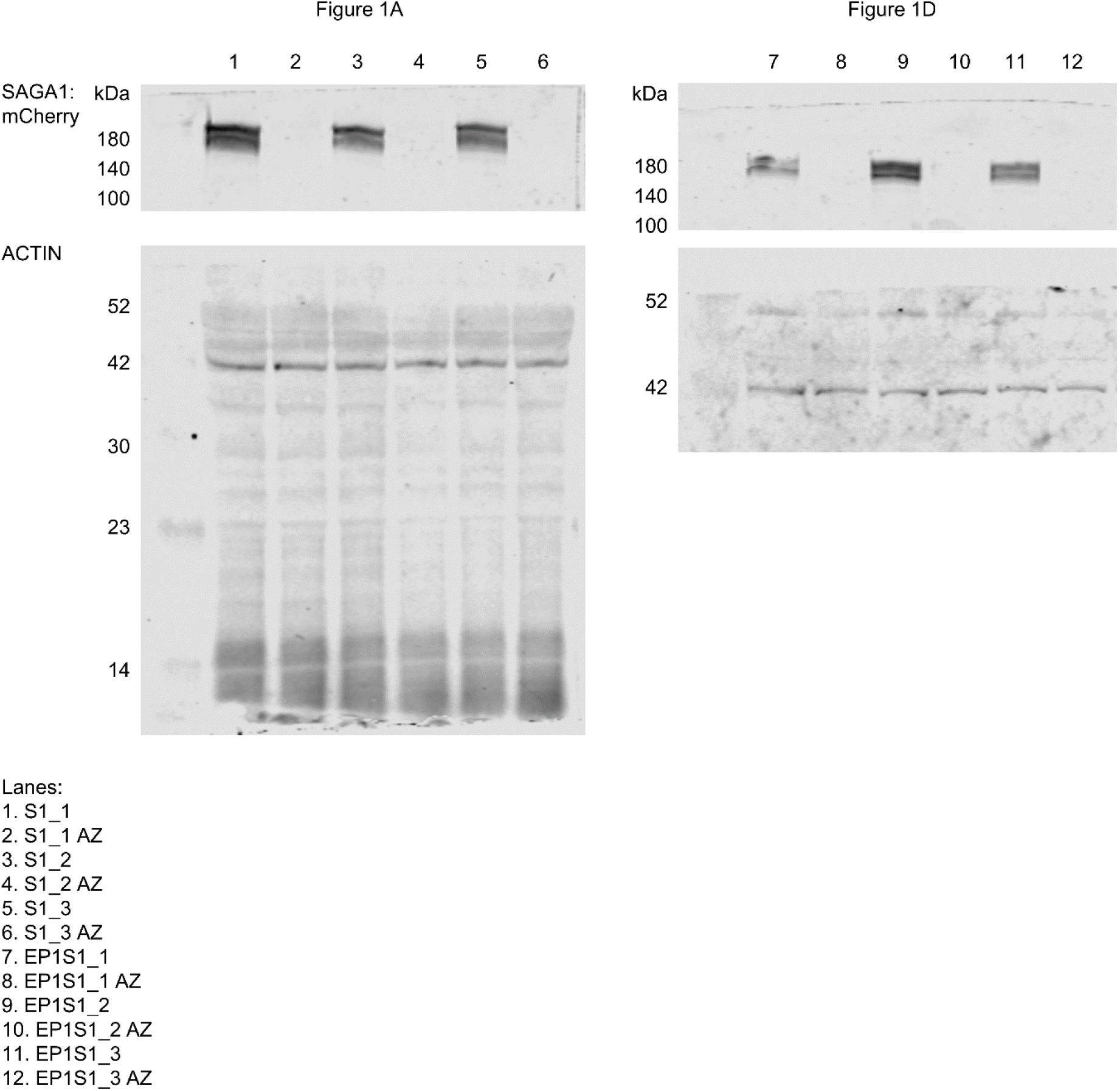
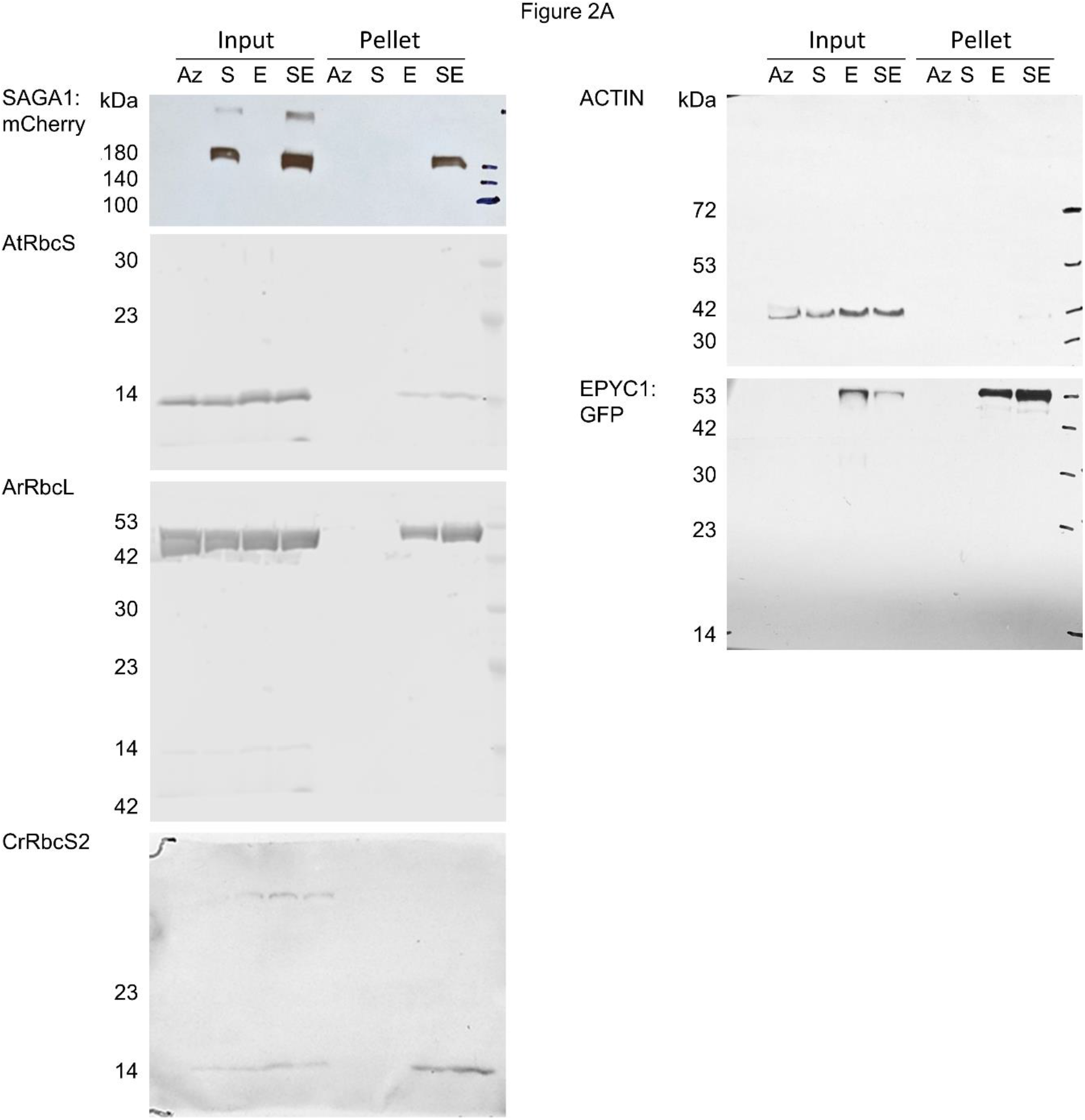
Sequence maps of plasmid vector used in this study. (*attached with submission*, ‘Figure S12 - Plasmid vector maps.zip’). See **Table S1** for a summary list of vectors.

**Figure S13.**
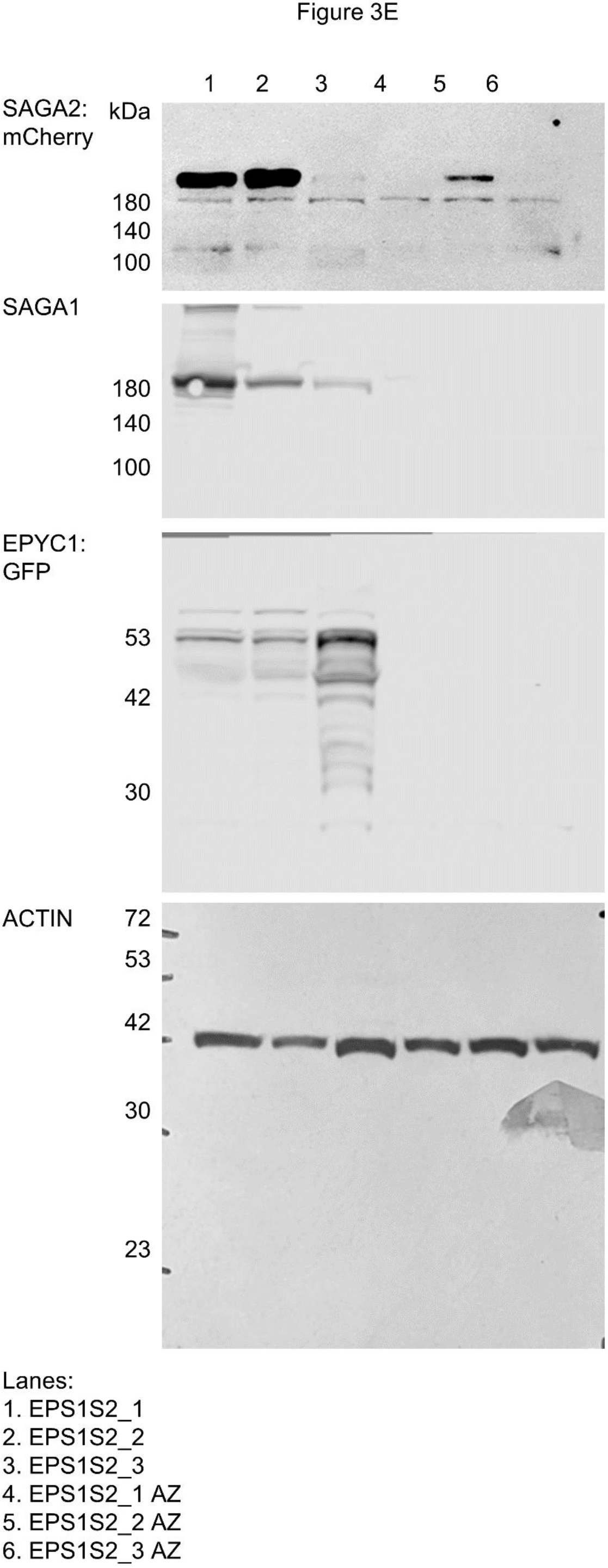
Uncropped immunoblot images from main figures.

**Figure S14.**
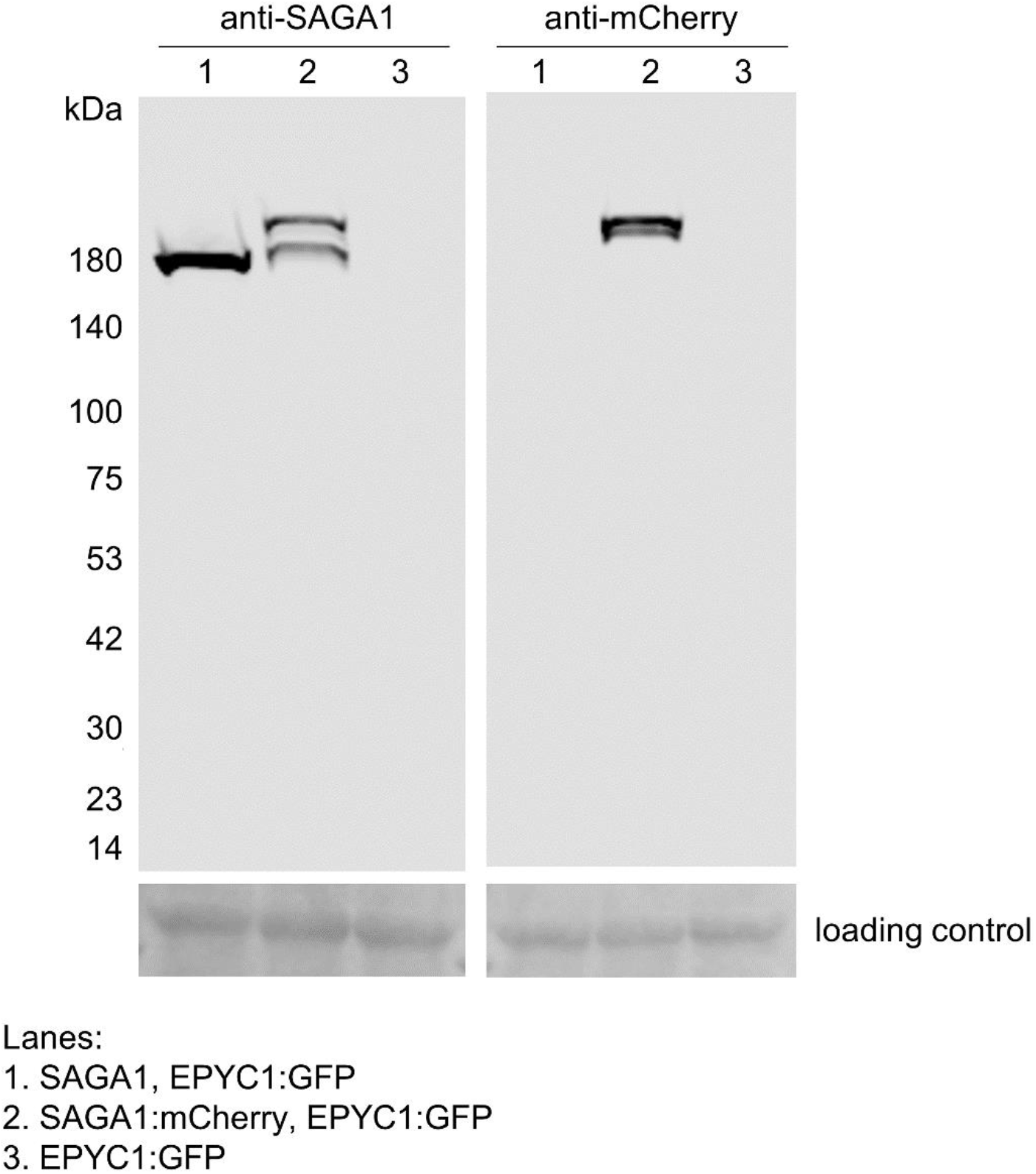
Immunoblots showing expression of SAGA1 and SAGA1:mCherry. Protein extracts from plants expression untagged SAGA1 and EPYC1:GFP (lane 1), SAGA1:mCherry and EPYC1:GFP (EP1S1, lane 2) were probed with either anti-SAGA1 or anti-mCherry antibodies. Plants expressing only EPYC1:GFP (lane 3, EP1) were included as a control. The higher bands in lane 2 correspond to SAGA1:mCherry, the lower bands correspond to untagged SAGA1. The presence of two bands in lane 2 for anti-SAGA1 indicates that mCherry is prone to cleavage (as seen in Fig. 1). The absence of free mCherry (27 kDa) indicates that cleaved mCherry is rapidly degraded.

**Table S1. List of plasmid vectors used in this study** (*attached with submission*, ‘Table S1 List of plasmid vectors.xlsx’). See **Fig. S12** for vector maps as .gb files.

**Table S2.** SBF-SEM staining protocol.

| Steps | Solution | Time | Temperature | Microwave watt | Vacuum |
| --- | --- | --- | --- | --- | --- |
| 1 | 1.5% Glutaraldehyde<br>2.5% Paraformaldehyde<br>5% Sucrose in 0.1 M NaCac | 2 x 14 min<br>(2min on/off cycles) | Room temperature | 100 | On |
| 2 | 0.1 M NaCac | 1 immediate.<br>Then 2 x 40 s. |  | 100 | On |
| 3 | 2% OsO<sub>4</sub> in 0.1 M NaCac | 2 x 14 min<br>(2min on/off cycles) |  | 100 | On |
| 4 | 2.5% K<sub>4</sub>[Fe(CN)<sub>6</sub>] · 3H<sub>2</sub>O<br>in 0.1 M NaCac | 2 x 14 min<br>(2 min on/off cycles) |  | 100 | On |
| 5 | Water wash | 1 immediate.<br>Then 2 x 40 s. |  | 100 | On |
| 6 | 1% thiocarbohydrazide<br>unbuffered | 2 x 14 min<br>(2 min on/off cycles) |  | 100 | On |
| 7 | Water wash | 1 immediate.<br>Then 2 x 40 s. |  | 100 | On |
| 8 | 2% OsO<sub>4</sub> aqueous | 2 x 14 minutes<br>(2 min on/off cycles) |  | 100 | On |
| 9 | Water wash | 1 immediate.<br>Then 2 x 40 s. |  | 100 | On |
| 10 | Dehydration series in<br>ethanol (25%, 50%,<br>75%, 3 x 100%) | 40 s each |  | 250 | Off |
| 11 | Infiltration series in<br>Durcupan:ethanol mixes<br>(25%, 50%, 75%, 90%,<br>2 x 100%) | 3 min each |  | 150 | On |
| 12 | 100% Durcupan | overnight |  | N/A | N/A |
| 13 | Polymerisation | 30 min | 60°C | 250 | Off |
| 14 | Polymerisation | 90 min | 100°C | 450 | Off |

**Table S3. Gene coding sequences used for Yeast Two-Hybrid assays** (*attached with submission*, ‘Table S3 Y2H sequences.xlsx’).

